# Dengue and Zika virus 5’-UTRs harbor IRES functions

**DOI:** 10.1101/557942

**Authors:** Yutong Song, JoAnn Mugavero, Charles B. Stauft, Eckard Wimmer

**Affiliations:** Department of Molecular Genetics and Microbiology, Stony Brook University, Stony Brook, NY, 11794; Current address: Codagenix Inc., Farmingdale, NY 11735; Co-founder of Codagenix Inc., Farmingdale, NY 11735

**Keywords:** dengue virus, Zika virus, cap-dependent translation, cap-independent translation, internal ribosome entry site: IRES

## Abstract

Members of *Flavivirus*, a genus of *Flaviviridae*, encompass numerous enveloped plus strand RNA viruses, of which globally dengue virus (DENV) is the leading cause of serious arthropod-borne disease. The genomes of DENV, just as those of yellow fever virus (YFV), West Nile fever virus (WNV), or Zika virus (ZIKV), control their translation by a 5’-terminal capping group. Three other genera of Flaviviridae are remarkable because their viruses use internal ribosomal entry sites (IRESs) to control translation and they are not arthropod transmitted. In 2006 E. Harris’ group published work suggesting that DENV RNA does not stringently need a cap for translation. They proposed that instead DENV translation is controlled by an interplay between 5’ and 3’ termini. Here we present evidence that the DENV or ZIKV 5’-untranslated regions (5’-UTRs) alone have IRES competence. This conclusion is based, first, on the observation that uncapped mono-cistronic mRNAs 5’ terminated with the DENV or ZIKV 5’-UTRs can efficiently direct translation of a reporter gene in BHK and C6/36 cells; second, that either 5’-UTR placed between two reporter genes can efficiently induce expression of the downstream gene in BHK but not in C6/36 cells. These experiments followed observations that uncapped DENV/ZIKV genomic transcripts, 5’ terminated with pppAN… or GpppAN…, can initiate infections of mammalian (BHK) or mosquito (C6/36) cells. IRES competence of the 5’-UTRs of DENV/ZIKV raises many open questions regarding the biology and control, as well as the evolution, of insect-borne flaviviruses.

**Importance:** Members of the genus *Flavivirus* of *Flaviviridae* are important human pathogens of great concern because they cause serious diseases, sometimes death, in human populations living in tropical, subtropical (dengue, DENV; Zika, ZIKV; yellow fever virus), or moderate climates (West Nile virus). Flaviviruses are known to control their translation by a cap-dependent mechanism. We have observed, however, that the uncapped genomes of DENV or ZIKV can initiate infection of mammalian and insect cells. We provide evidence that the short 5’ untranslated region (5’-UTR) of DENV or ZIKV genomes can fulfill the function of an internal ribosomal entry site (IRES). This strategy frees these organisms from the cap-dependent mechanism of gene expression at an as yet unknown stage of proliferation. The data raise new questions about the biology and evolution of flaviviruses, possibly leading to new controls of flavivirus disease.

## Introduction

The 1988 discovery of the “Internal Ribosomal Entry Site (IRES)” in our and in Sonenberg’s laboratories changed research on translational control in eukaryotic systems (1, 2). At the time cap-dependent translation in eukaryotic cells had been elevated to a dogma, which was suddenly punctured by an alternative mechanism.

IRESs were originally discovered in picornavirus genomes (that function as mRNA) where they are long, highly structured RNA segment up to 450 nt long (1, 2). The IRES in the intragenic region (IGR) of the insect pathogen cricket paralysis virus (CrPV), on the other hand, is only 189 nt long (3). IRESs have also been discovered to function in the expression of cellular genes and, again, they are of different structures and sizes (4). These properties make it difficult to identify IRESs by bioinformatics. The observation of a minute viral IRES (96 or 107 nt long) described here is intriguing.

Members of the family *Flaviviridae* comprise a large group of pathogenic viruses, of which many cause severe diseases in millions of humans globally (5). These viruses are enveloped, +ssRNA viruses with genomes approximately 11 kb in length that are not 3’ polyadenylated (6). They encode a single polypeptide, the polyprotein, consisting of an array of related structural and non-structural proteins (5).

It is noteworthy that *Flaviviridae* have evolved into two distinct groups with profound differences in their life cycle, especially in the strategy to control translation. *Group One* comprises member viruses of the genera *Hepacivirus* (hepatitis C virus), *Pestivirus* (bovine viral diarrhea virus), and *Pegivirus* (GB virus). All of them are blood-borne or are transmitted by contact. These viruses control their translation strictly through internal ribosomal entry sites (IRESs) (6). *Group Two* belongs to the genus *Flavivirus* and contains a very large number of species most of which are transmitted by, and can replicate in, insects or acarine species (“arboviruses”). The best-known human/primate flavivirus is dengue virus (DENV), a health threat to 2.5 billion humans in tropical and subtropical climates. More recently, Zika virus (ZIKV), an agent closely related to DENV, has emerged as a new dangerous flavivirus that co-circulates with DENV in tropical and sub-tropical climates. As an accepted rule, each member of the genus *Flavivirus* controls its translation by a “cap-dependent” (m^7^GpppANNN…) mechanism (6).

However, a previous study suggested that DENV can initiate protein synthesis by using a cap-independent manner through interaction between its 5’- and 3’-UTR (7). Experiments have led the authors to conclude that no IRES-like function is involved. Stimulated by results from E. Harris’ laboratory (7) we tested whether our purified, whole length, non-capped transcript RNAs of DENV and ZIKV cDNAs can infect mammalian or mosquito cells by transfection. Surprisingly, the non-capped flavivirus genomes readily infected mammalian (BHK, Vero) or mosquito cells (C6/36) and produced virus in high titers. Translation is the initial step of replication in infections by naked +stranded RNA genomes. In our case, there are various hypothesis to explain what could lead to viral protein synthesis directed by the uncapped RNA transcripts: translation of uncapped genomes, translation of genomes that were capped in the cytoplasm, or translation controlled by the genomic 5’UTRs that harbor IRES competence. These possibilities will be discussed later. We opted to focus on the third hypothesis that predicts cap-independent initiation of translation through IRES function of the small 5’-UTRs.

Our experiments followed established strategies: control of translation in mono-cistronic mRNAs with *Gaussia* luciferase (*Gluc*) as reporter followed by experiments in di-cistronic mRNA. The di-cistronc mRNAs consisted of the firefly luciferase (*Fluc*) gene and the gene for *Gluc* while the DENV or ZIKV 5’UTRs were introduced intergenically between the *Fluc* and *Gluc* genes (1, 2). The results of the experiments reported here allow us to suggest that the short nucleotide sequences of the 5’-UTRs of DENV (96 nt) and ZIKV (107 nt) are competent to serve as very small IRESs directing initiation of translation.

## Results

### The DENV genomic RNAs can be translated cap-dependently and cap-independently in both mammalian and mosquito cells

In all of our experiments described here, we have used our synthetic DENV type 2 clone based on wt DENV2 strain 16681 (8). This plasmid was equipped with a T7 phi2.5 promoter that synthesizes RNA starting with adenosine, the first base (following the cap) of the wt DENV genome (9, 10). Using phage T7 RNA polymerase we synthesized transcript RNAs and modified the transcripts with one of three different 5’-termini: cap-unrelated (pppAN…), a non-functional cap (GpppAN…, non-methylated), or a functional cap (m^7^GpppAN…). Transfection of individual transcripts either into mammalian (BHK) or into mosquito (C6/36) cells produced virus at the end point of experiments roughly equal to that of the transcripts with the functional m^7^G cap (Fig. 1). As we have observed previously (8), infectious virus yields were always slightly higher in C6/36 cells than in BHK cells. This observation suggests that DENV infection is more productive in mosquito cells compared to mammalian cells.

**Fig. 1.**
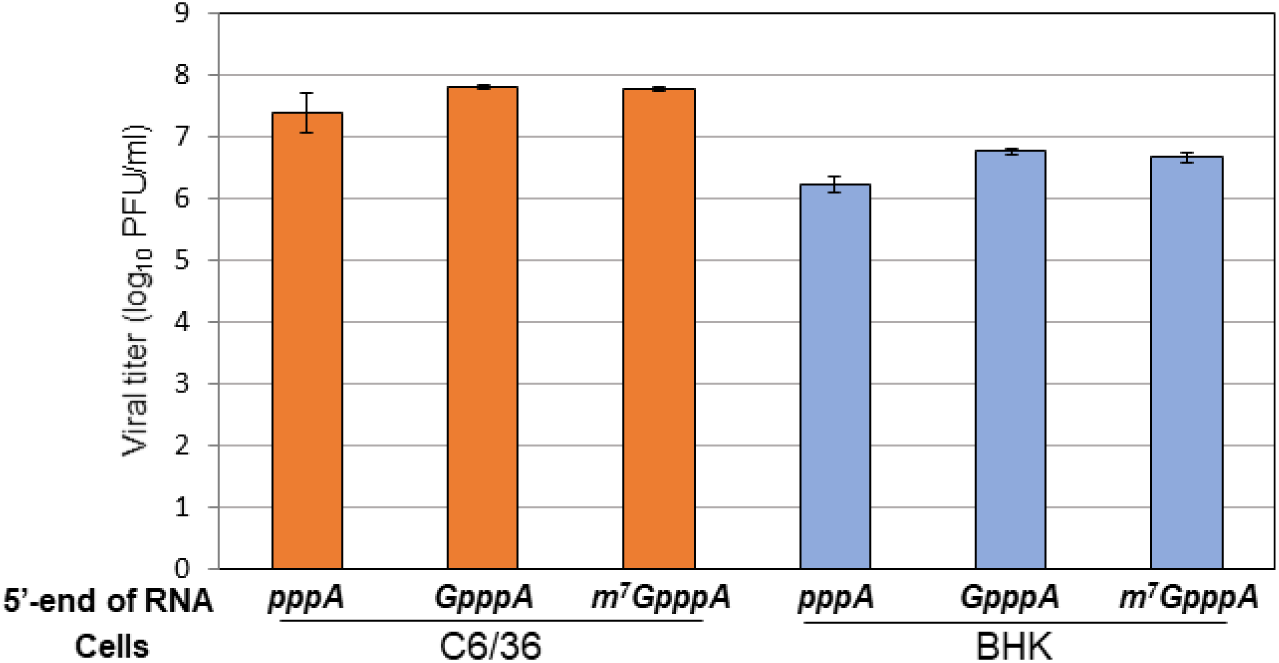
A non-modified 5’-end DENV genome RNA can produce infectious viral particles in both mammalian and mosquito cells. BHK cells and C6/36 cells seeded on a 12-well plate were transfected with equal amounts of three types of in-vitro transcripts: (1) a non-modified 5’pppA-terminated RNA, (2) an unmethylated 5’ GpppA-terminated RNA, and (3) a standard m^7^GpppA-capped RNA generated from the synthetic DENV2 infectious clone (DENV2^Syn^, (8)). The cell supernatants were harvested at 5 days post transfection (CPE can be detected in BHK cells, but not in C6/36 cells). Infectious viral titer (PFU/ml) was determined by plaque assay using BHK cells. The data are expressed as an average of two experiments. Error bars indicate standard deviations.

Both the m^7^GpppA-capped and unmethylated GpppA-capped DENV transcripts produced similar viral titers on BHK cells post transfection (Fig. 1). Therefore, we carried out a detailed assay of infectivity with the two uncapped pppAN- and GpppAN-DENV genome transcripts wondering if we would find major differences in specific infectivity of the *in vitro* RNAs. Samples were harvested daily (up to day 5) from medium and used for focus forming assays in Vero cells (see Materials and Methods). The result showed that the pppA-DENV2 genome RNA produced viral titers that were 2-3 log_10_ lower than those obtained with the unmethylated GpppA-5’modified RNAs (Fig. S1). This observation is in accordance with a previous report obtained with transcripts of yellow fever virus cDNA (11). We assume that the viruses harvested at the end of the incubation contain genomes with DENV-specific capping groups. The reason is that the viral replication machinery, that was newly assembled in the course of the replication cycle, will provide newly synthesized genomes with the 5’ terminal modification (12-14). Experiments to test this hypothesis are currently in progress.

As was mentioned in the Introduction, there are different possibilities to explain the infectivity of uncapped DENV transcripts (see Discussion). In the following we report our experiments to test the DENV and ZIKV 5’ UTRs for IRES competence.

### Translation of mono-cistronic mRNAs under the control of DENV 5’-UTR variants in mammalian cells

To test whether the uncapped transcripts of the DENV cDNA harbor an activity that allows cap-independent translation we designed mono-cistronic mRNAs 5’ terminated with either pppAN, GpppAN or m^7^GpppAN, followed by the 96-nt-long DENV 5’-UTR sequence (***D5***; Fig. 2) and the *Gluc* ORF (15). We chose two different 3’ termini, the DENV 3’-UTR (***D3***) because of its possible role in DENV translation (16) or the polyadenylated 3’-UTR (P) of poliovirus because of evidence that 3’-terminal poly(A) enhances IRES-mediated initiation of translation (17). Furthermore, the 3’ termini may also contribute to mRNA stability. These constructs are abbreviated as, for example, pppA-***D5***G***D3*** or pppA-***D5***GP, respectively (Fig. 2A). Altogether we prepared 12 different transcripts carrying different termini (detailed in the Table of Fig. 2A). Since the DENV 5’ cyclization sequence (5’-CS), located in the 5’ part of core protein-encoding sequences, has not been proven to be necessary for efficient translation (18), we did not retain the 5’-CS in these mono-cistronic constructs. To standardize the translation experiment, we determined that ∼400 ng of DENV-*Gluc* mRNA [m^7^G-***D5***G***D3***] yielded an optimized signal in transfected mammalian cells (Fig. S2). About 400 ng mono-cistronic mRNA were therefore selected for all subsequent intracellular translation assays. RNA transcripts were individually transfected into BHK cell monolayers using Lipofectamine®2000, and *Gluc* activities were measured at different time-points post transfection (as indicated; for details, see Materials and Methods). The results showed that the non-methylated (GpppA) RNAs generated *Gluc* expression levels surpassing the signal of the m^7^G capped mRNA (Fig. 2B), a result suggesting that in this experiment the classical m^7^G cap is not required for efficient translation. We expected that the DENV 3’-UTR may contribute to *Gluc* expression particularly with pppA-***D5***G***D3*** mRNA (compare Fig. 2B with 2C) and, indeed, we observed that the pppA-***D5***G***D3*** reporter RNA produced relatively high *Gluc* activity (pppA-***D5***G***D3***, Fig. 2B) but low *Gluc* with pppG-GP (Fig. 2D). We note that in these experiments the unrelated 3’-untranslated sequences “minus poly(A)” or “plus poly(A)” were of little, if any, effect on translation of mRNAs without a functional m^7^G cap (GP, Fig. 2D and 2E). The qRT-PCR results showed that the differences in translational efficiencies are not due to RNA instability of the different reporter RNAs isolated from the transfected cells (Fig. S3).

**Fig. 2.**
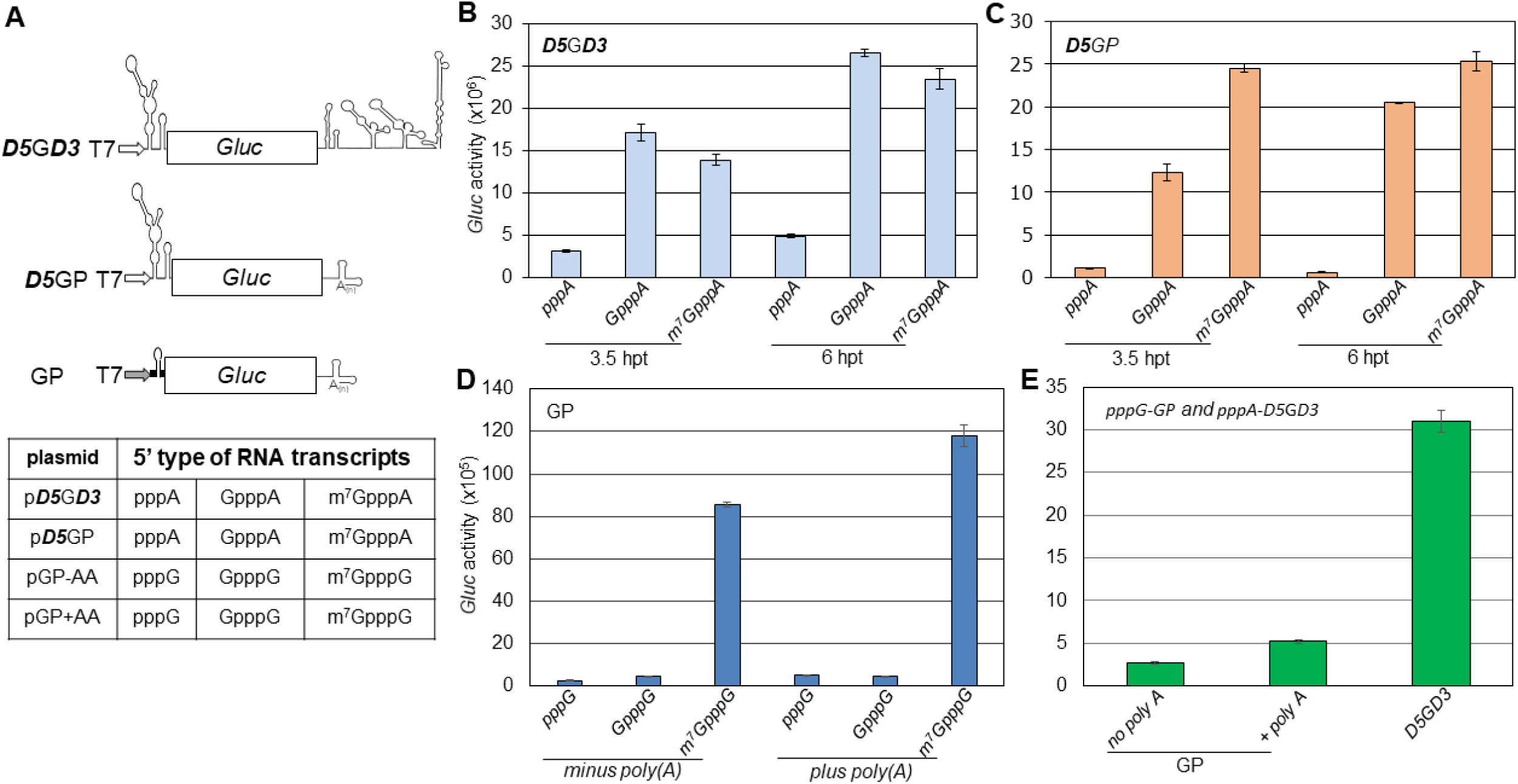
Translation assay directed by the DENV 5’-UTR with different 5’-modifications in BHK cells. **(A)** Diagram of DENV mono-cistronic reporter constructs (details in Materials and Methods). RNA transcripts were generated from a modified/standard T7 promoter. Different types (as shown below X-axis) of RNA transcripts: ***D5***G***D3* (B)**, ***D5***GP **(C)**, and GP **(D)** were generated from a modified (open arrow, for **B** and **C**)/standard (solid arrow, for **D**) T7 promoter, and then transfected into BHK monolayer cells seeded on a 12-well plate. A certain volume of the medium was harvested for measurement of luciferase activity at different time-points **(B**, **C)** or at 3.5 hours post transfection (hpt) **(D**, **E)**. **(E)** Comparison of *Gluc* expression from non-capped RNA transcripts of ***D5***G***D3*** and GP. Means of four independent experiments are plotted ± SEM. *Gluc, Gaussia* luciferase reporter gene.

These results strongly suggest that the genome of DENV harbors in its 5’-UTR a structure that, independently of a cap structure or the DENV 3’-UTR, is capable of initiating translation.

### In mammalian cells the DENV 5’-UTR activates internal ribosomal entry in di-cistronic mRNAs

Based on the results with mono-cistronic mRNAs we tested a possible IRES function of the DENV 5’-UTR in di-cistronic reporter mRNAs. These mRNAs contained 5’-terminal the *Fluc* and 3’-terminal the *Gluc* genes with the 96 nt of the DENV 5’-UTR placed between two reporter genes. These mRNAs, designated F***D5***G***D3*** (*Fluc*-DENV2 5’-UTR-*Gluc*-DENV2 3’-UTR) and F***D5***GP (*Fluc*-DENV2 5’-UTR-*Gluc*-PV 3’-UTR), allowed us to assess a possible IRES competence of the 5’-UTR and a contribution of the DENV 3’-UTR to translation initiation of *Gluc* (Fig. 3B). Two additional di-cistronic mRNAs, F***H5***GP and F***ΔH5***GP, carry the entire HCV IRES, or a deletion version thereof (Fig. 3B) and were used as positive and negative controls, respectively. The 5’ termini of these mRNAs (bold lines) are an uncapped, DENV-unrelated sequence (n=41). Importantly, this sequence has the propensity to form a stable hairpin to reduce fortuitous initiation of translation, thereby avoiding an undesired high background of the *Fluc* signal (Fig. 3A).

**Fig. 3.**
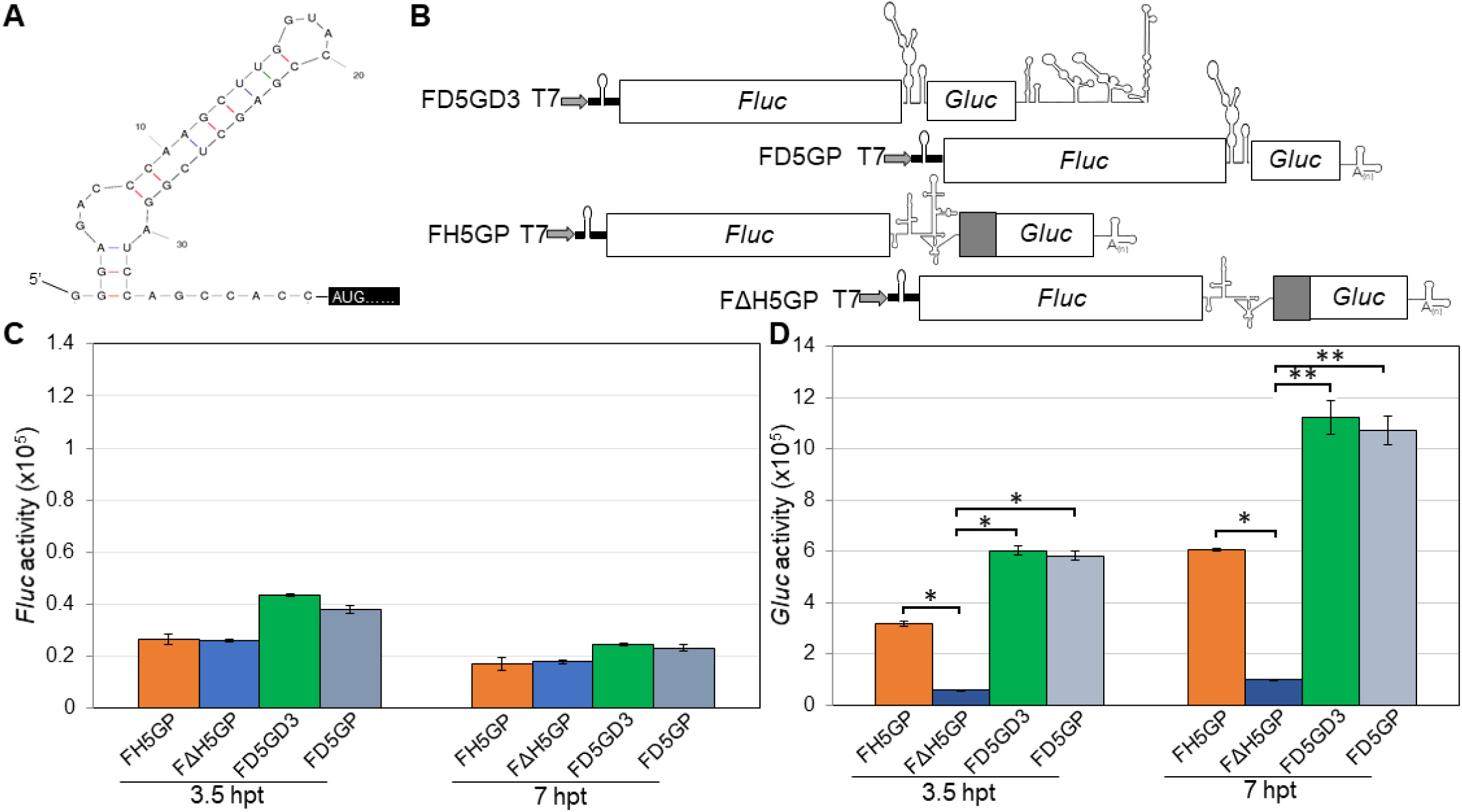
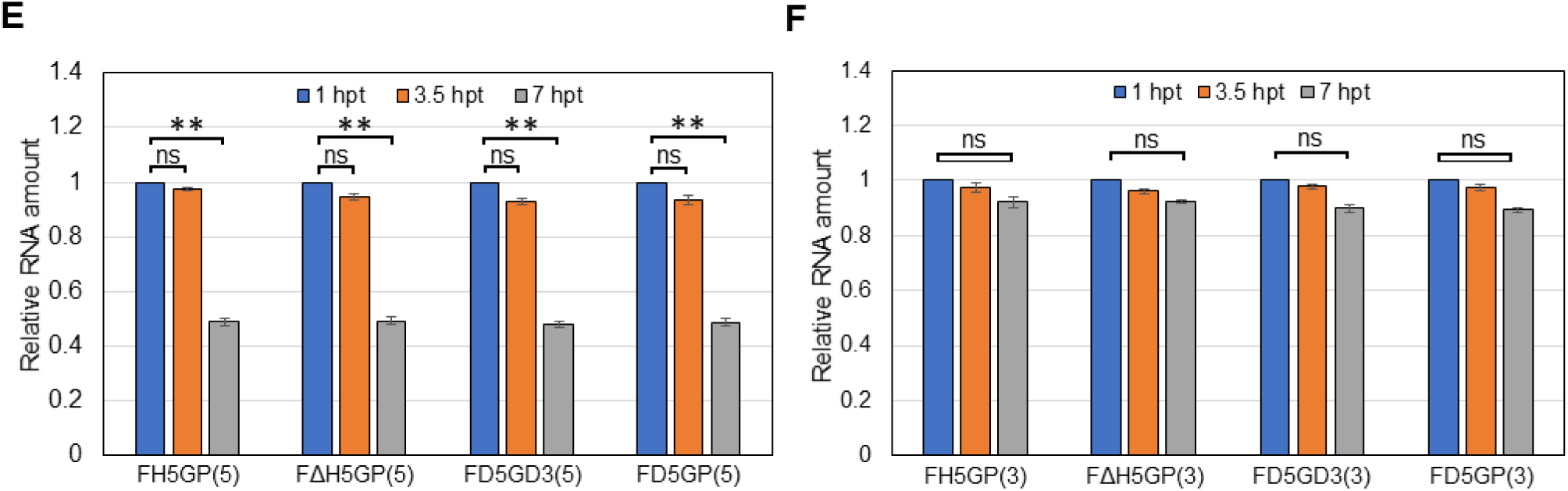
The DENV2 5’-UTR confers IRES activity in di-cistronic mRNAs. (**A**) Sequence and predicted secondary structure of 5’ terminal UTR of all di-cistronic constructs in this study are shown. ΔG = −13.8 calculated by RNA mfold. **(B)** Schematic diagram of di-cistronic reporter constructs F*D5*G***D3***, F***D5***GP, F***H5***GP, and F***ΔH5***GP. **(C)** Translation efficiency of 5’ non-capped *Fluc* reporter RNAs. F***H5***GP and F***ΔH5***GP reporter RNAs were included as positive and negative control, respectively. **(D)** *Gluc* expression directed by internal initiation of either the HCV IRES or the DENV2 5’-UTR. *Fluc* and *Gluc* activities were measured at different time-points post transfection as indicated. The means of three independent experiments are plotted ± SEM. *Fluc*, firefly luciferase reporter gene. The P values were determined by comparing to the negative control (F***ΔH5***GP) using two-tailed t-tests at the indicated time point. * and ** indicate P<0.05 and P<0.01, respectively. **(E, F) Relative amount of four di-cistronic RNA transcripts in transfected cells analyzed by qRT-PCR.** Total RNA was extracted and isolated by *Trizol*^®^ reagent from cells transfected with RNA transcripts at different time-points post transfection. Relative RNA levels were measured by qRT-PCR with either 5’-oligo primer pairs **(E)** or 3’-oligo primer pairs **(F)** and calibrated by the RNA amount at 1 hpt, respectively. Final percentage of RNA amounts were normalized by GAPDH. The means of three independent experiments are plotted ± SEM. A significant difference by two-tailed t-tests compared to RNA amount (at 1 hpt) is indicated by ** (P<0.01). ns, not significant.

Individual transfection of these transcripts into the BHK cells revealed weak upstream *Fluc* expression in all cases but robust downstream *Gluc* expression as controlled by the inter-cistronic DENV 5’-UTR (Fig. 3). In fact, downstream expression of *Gluc* activated by the DENV 5’-UTR was nearly twice as strong as the activation by the HCV IRES. We note that the nature of the 3’-termini (DENV 3’-UTR, F***D5***G***D3***; or PV 3’-UTR, F***D5***GP) of the di-cistronic mRNAs played at best a minor role in these experiments (Fig. 3D). This result supports our conclusion that the 3’-UTR is unlikely to play a critical role, if any, in upstream *Gluc* expression. Relative *Gluc* expression levels normalized to *Fluc* activity are shown in Fig. S4A. Our qRT-PCR results showed that the difference in translational efficiencies is unrelated to RNA stabilities of the reporter RNAs isolated from the transfected cells (Figs. 3E and 3F).

In addition, the expression of *Fluc* and *Gluc* from the *same but capped* dicistronic mRNA led to a very high *Fluc* signal (Fig. S4B), as expected. In contrast, the *Gluc* signal (Fig. S4C) from the same capped dicistronic mRNA was very similar as the signal obtained from the corresponding uncapped dicistronic mRNA analyzed earlier (Fig. 3D). This suggests very little influence of ORF1 expression on ORF2 expression under the conditions of our experiment.

We note that the findings reported here do not conform with established translation initiation. We conclude that translation in these di-cistronic mRNAs is mediated by an RNA segment with “internal ribosome entry” competence. The DENV2 5’-UTR (96 nt) therefore can function as a minute IRES element that induces robust translation of a downstream gene but only a very weak translation of an upstream gene.

### The DENV 3’-UTR does not play a critical role in the non-canonical translation initiation in mammalian cells

Following the strategy employed for the DENV 5’-UTR we placed the long 3’-UTR (451 nt) into the intergenic region of the di-cistronic mRNAs (F***D3***GP, Fig. S5A) and tested the activity of the reporter genes after transfection into BHK cells. The 3’-UTR, however, did not initiate significantly non-canonical translation of either the upstream *Fluc* or the downstream *Gluc* genes (Figs. S5B and S5C). In contrast. the IRES of encephalomyocarditis virus (EMCV), which was used as an intergenic control (F***E***GP), caused very strong downstream activation of the *Gluc* gene (Fig. S5C).

The untranslated regions of flavivirus genomes engage in complex interactions with distant upstream genomic sequences, which have been recognized as crucial for viral proliferation (19, 20). Although the DENV 5’-UTR that we have so far analyzed lacks some of the possible binding sites for the 3’-UTR, we constructed di-cistronic mRNAs with both 5’- and 3’-UTRs between the two genes in either orientation (F***D53***GP and F***D35***GP). This allowed us to test whether any interaction takes place between the terminal RNA segments that might yield translation cooperativity during this non-canonical translation initiation. The results suggested that there was some stimulation of downstream translation under the control of F***D35***GP RNA but the activation of *Gluc* was weak.

### Activation of translation by the DENV 5’-UTR in mono-cistronic mRNAs in mammalian and mosquito cells

Being an arbovirus, DENV can infect and replicate in mosquitoes. We therefore expected that in C6/36 mosquito cells the DENV 5’-UTR would initiate cap independent translation of mono-cistronic mRNA as described for experiments in BHK cells (Fig. 2). We used ***D5***G***D3*** mRNAs with different 5’ ends (pppA-, GpppA-, m^7^GpppA-) as described in Fig. 4 and, in addition, mono-cistronic mRNA with the HCV IRES preceding the *Gluc*, as control (***H5***G***H3***, Fig. 4A). Just as in BHK cells, the non-capped pppA-***D5***G***D3*** RNA produced a low *Gluc* signal in C6/36 cells, an observation that we will address later. mRNAs terminated with 5’ GpppA or m^7^GpppA, however, produced a robust *Gluc* signal (Fig. 4B). In contrast, the ***H5***G***H3*** RNA, containing the HCV IRES upstream of *Gluc* failed to activate translation of *Gluc* in the mosquito cells (Fig. 4C). This supports a recent report that the IRES of a virus of the genus *Hepacivirus* of *Flaviviridae* is non-functional in insect cells (21).

**Fig. 4.**
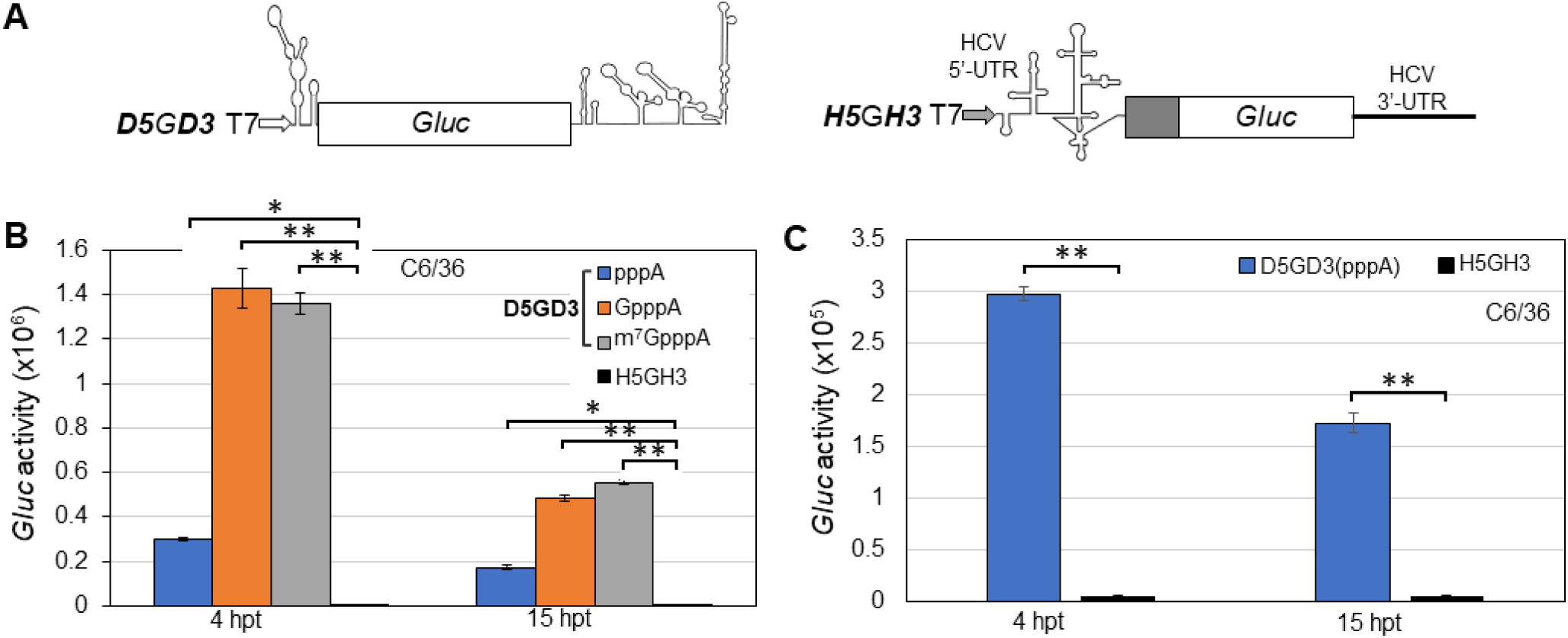
Translation of mono-cistronic RNAs containing the DENV2 5’-UTR with different 5’-modifications in C6/36 cells. **(A)** Diagram of DENV and HCV mono-cistronic reporter constructs ***D5***G***D3*** and ***H5***G***H3***. **(B)** Different types of mono-cistronic ***D5***G***D3*** RNA transcripts and one type of ***H5***G***H3*** reporter RNA were transfected into C6/36 cells, and the *Gluc* activities were measured at different time-points post transfection as indicated. **(C)** Comparison of *Gluc* expression from only non-capped RNA transcripts of ***D5***G***D3*** and ***H5***G***H3*** at 3.5 hpt. The means of three independent experiments are plotted ± SEM. The P values were determined by comparing to the negative control (***H5***G***H3***) using two-tailed t-tests at the indicated time point. * and ** indicate P<0.05 and P<0.01, respectively.

### In mosquito cells, the DENV 5’-UTR placed between *Fluc* and *Gluc* in di-cistronic mRNA is unable to activate translation of *Gluc*

Considering the results of mono-cistronic mRNAs in C6/36 mosquito cells we expected that the DENV 5’-UTR too would initiate translation of a downstream gene in di-cistronic mRNAs in C6/36 mosquito cells. To our surprise this is not the case.

Specifically, we tested several different non-capped di-cistronic mRNAs. These include F***D5***G***D3*** and F***H5***GP, and the new constructs F***E***GP harboring the IRES of encephalomyocarditis virus (EMCV IRES), and F***Cr***GP harboring the small IRES of the intergenic region (IGR) of CrPV, an insect virus, respectively (Fig. 5A). The *Fluc* signals with any of these di-cistronic mRNAs were barely detectable (Fig. 5B). Unexpectedly, the downstream *Gluc* signals produced by the F***D5***G***D3***, F***H5***GP, or F***E***GP mRNAs were also extremely weak (Fig. 5C). The exception was, not surprisingly, the robust signal induced by F***Cr***GP mRNA, carrying the small IRES of an insect virus (Fig. 5C). Our qRT-PCR results showed that the difference in translational efficiencies is not due to RNA stabilities of the reporter RNAs isolated from the transfected cells (data not shown).

**Fig. 5.**
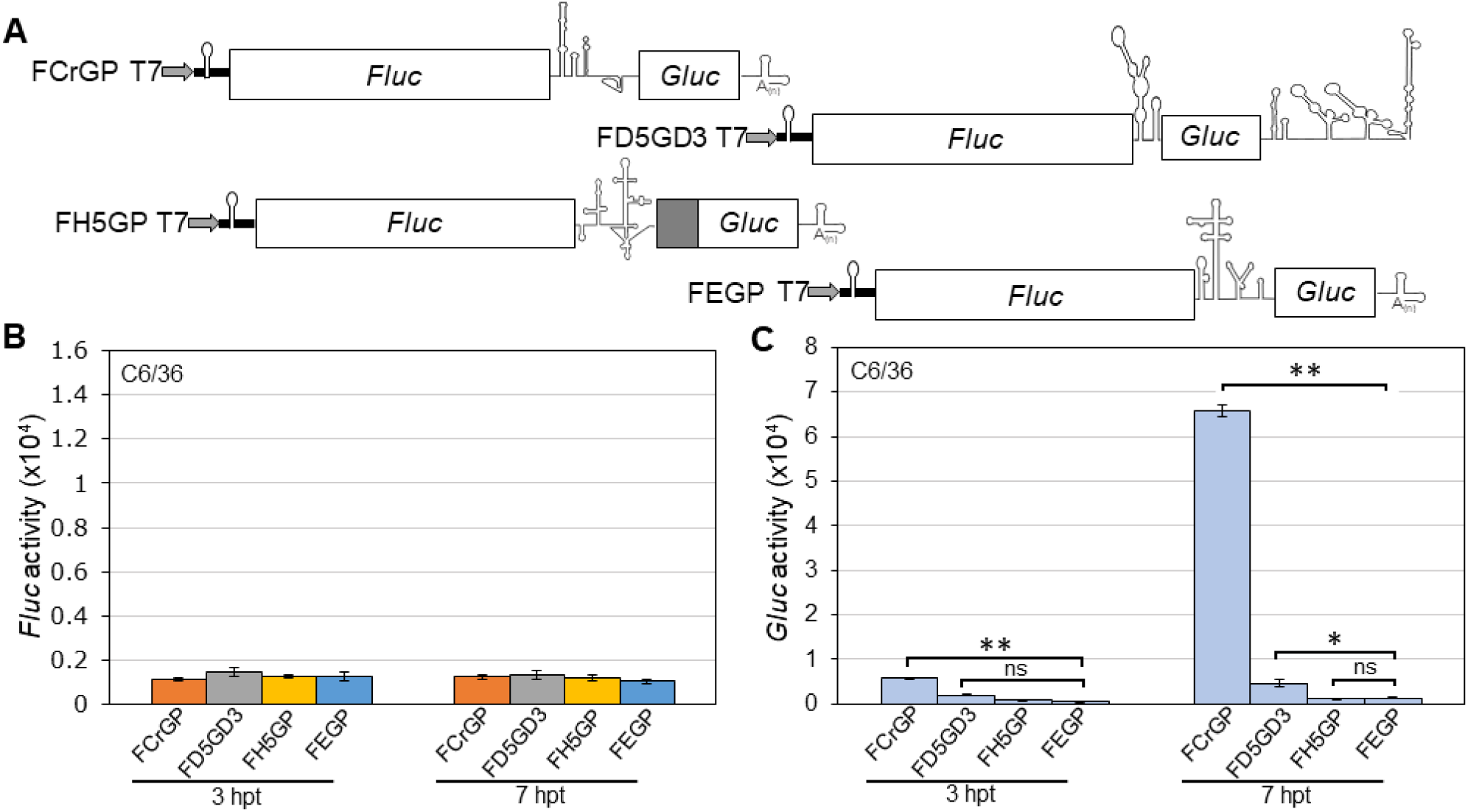
Translation assay of di-cistronic constructs containing the DENV 5’-UTR in C6/36 cells. **(A)** Schematic diagram of new di-cistronic reporter constructs F***E***GP and F***Cr***GP. **(B)** Translation efficiency of 5’ non-capped di-cistronic reporter RNAs. **(C)** *Gluc* expression directed by internal initiation of either the IGR CrPV IRES, EMCV IRES, HCV IRES, or the DENV2 5’-UTR. *Fluc* and *Gluc* activities were measured at different time-points post transfection as indicated. The means of three independent experiments are plotted ± SEM. The P values were determined by comparing to the negative control (F***H5***GP) using two-tailed t-tests at the indicated time point. * and ** indicate P<0.05 and P<0.01, respectively. ns, not significant.

The poor performance of the EMCV and HCV IRESs in mosquito cells is consistent with previous reports (21, 22). It came as surprise, however, that the DENV2 5’-UTR that had shown robust function in mono-cistronic mRNAs in insect cells, was nearly inactive in these cells when placed between the *Flul* and *Gluc* genes. We speculate that one (or more) specific protein factor(s) may be required in mosquito cells for the minute IRES of the DENV to function and that these factors are either absent or in too low quantities in C6/36 cells to activate translation.

### The 5’-UTR of the ZIKV genome reveals IRES competence similar to that of the DENV 5’-UTR

Many features of the molecular biology of ZIKV are closely related to DENV. Since the putative secondary structures of the respective 5’-UTRs share similar folding and stability (Fig. S7), we suspected that the ZIKV 5’-UTR may also have IRES competence. We tested this with experiments very similar to those with the DENV 5’-UTR. Using infectious ZIKV cDNA (FSS13025, GenBank number KU955593.1), kindly provided by Dr. Pei-Yong Shi (Galveston, TX), we first tested the infectivity of full-length ZIKV transcript RNAs, terminated with pppAN, GpppAN, or m^7^GpppAN in mammalian (Vero) and mosquito (C6/36) cells. Vero cells were chosen for ZIKV because plaque assays in these cells yielded superior results when compared to those with BHK cells. At the end point of the experiments all uncapped genome variants produced infectious virus in yields roughly equal to that of the genome RNA carrying a functional m^7^G cap (Fig. 6). Possible mechanisms leading to this result will be discussed in the Discussion.

**Fig. 6.**
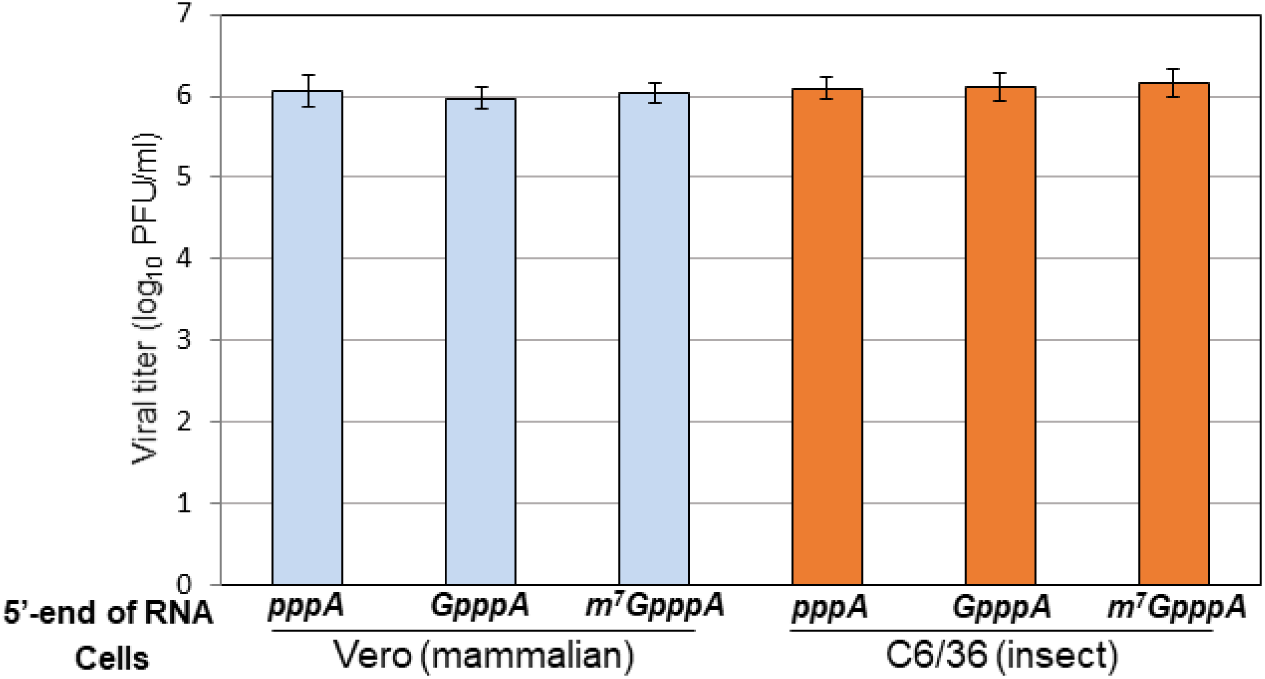
Non-capped ZIKV genome RNAs can produce infectious viral particles in both mammalian and mosquito cells. Vero cells and C6/36 cells seeded on a 12-well plate were transfected with equal amounts of three types of in-vitro transcripts: (1) a non-modified 5’pppA-terminated RNA, (2) a 5’ GpppA-terminated RNA, and (3) a standard m^7^GpppA-capped RNA generated from the pFLZIKV infectious clone (43). The cell supernatants were harvested at 5 days post transfection (CPE can be detected in Vero cells, but not in C6/36 cells). Infectious viral titer (PFU/ml) was determined by plaque assay using Vero cells. The data are expressed as an average of two experiments. Error bars indicate standard deviations.

We generated mono-cistronic mRNA ***Z5***G***Z3*** (Fig. 7A) and di-cistronic mRNA F***Z5***G***Z3*** (Fig. 8A) and tested these mRNAs in BHK cells. Comparison of ***Z5***G***Z3*** mRNA with ***D5***G***D3*** showed active translation of *Gluc* in both cases. We note that the signal from ***Z5***G***Z3*** surpassed that of the ***D5***G***D3*** RNA (Fig. 7B).

**Fig. 7.**
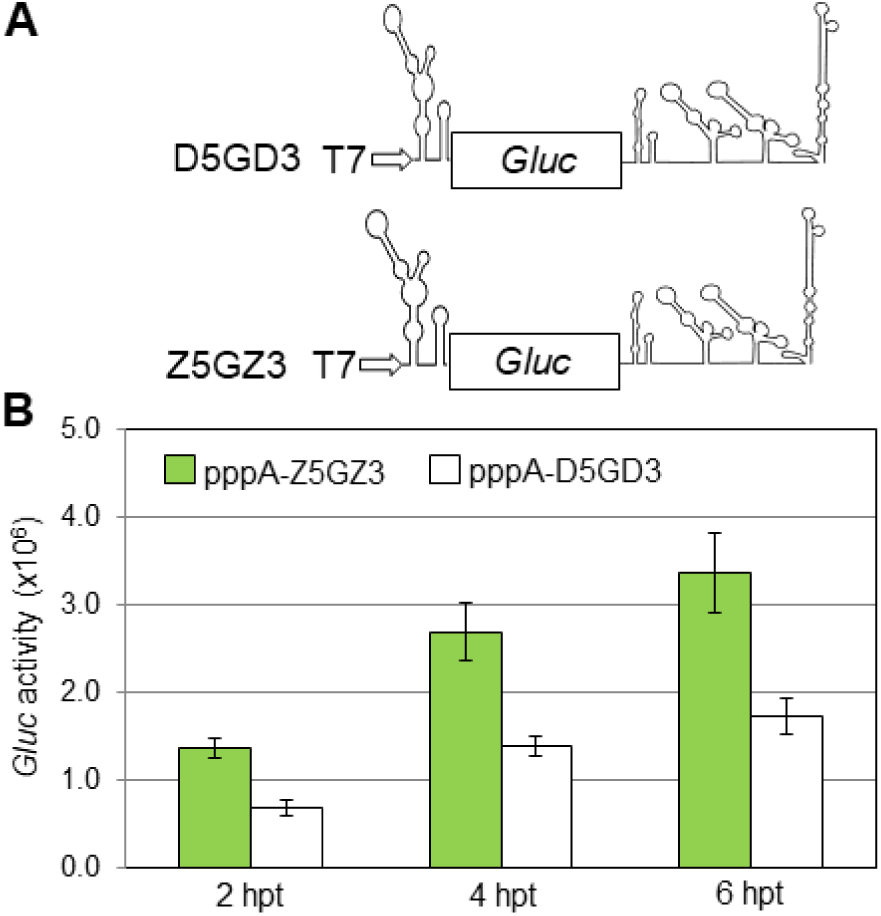
Translation assay of mono-cistronic constructs containing the ZIKV 5’-UTR in mammalian BHK cells. **(A)** Schematic diagram of ***D5***G***D3*** and a new mono-cistronic reporter construct ***Z5***G***Z3***. **(B)** *Gluc* expression level directed by the mono-cistronic pppA-***D5***G***D3*** and pppA-***Z5***G***Z3*** RNAs. The means of four independent experiments are plotted ± SEM.

**Fig. 8.**
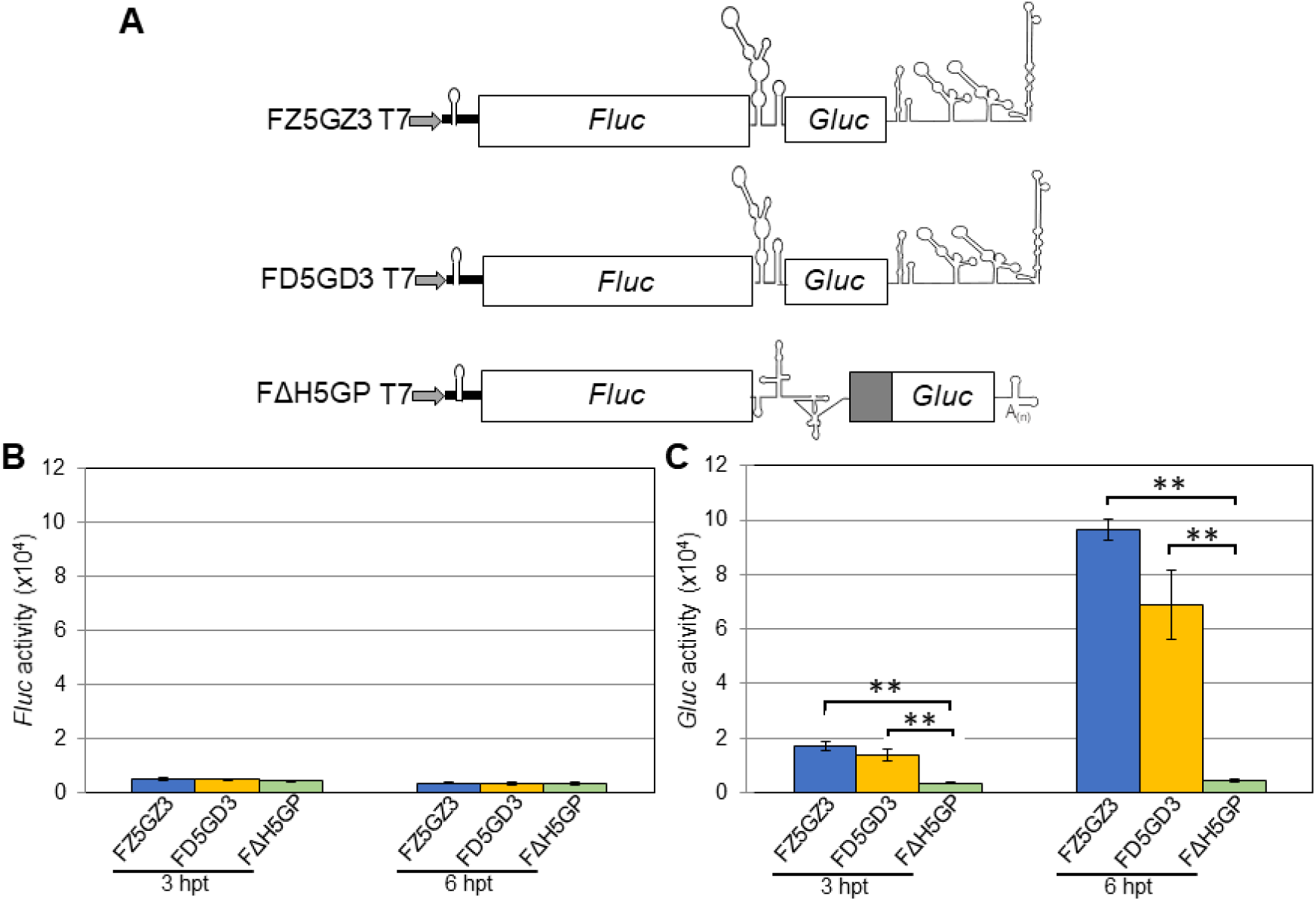
The ZIKV 5’-UTR shows a novel mechanism of translation initiation that is m^7^G-cap-independent, but IRES-dependent in mammalian BHK cells. **(A)** Diagram of F***D5***G***D3***, F***ΔH5***GP, and a new ZIKV di-cistronic reporter constructs F***Z5***G***Z3***. **(B)** Translation efficiency of 5’ non-capped di-cistronic reporter RNAs. **(C)** *Gluc* expression directed by internal initiation of either the HCV IRES or the DENV2 5’-UTR. *Fluc* and *Gluc* activities were measured at different time-points post transfection as indicated. The means of four independent experiments are plotted ± SEM. The P values were determined by comparing to the negative control (F***ΔH5***GP) using two-tailed t-tests at the indicated time point. ** indicates P<0.01.

We then tested three di-cistronic reporter mRNAs, F***Z5***G***Z3*** (ZIKV 5’- and 3’-UTRs), F***D5***G***D3*** (DENV 5’- and 3’-UTRs) and F***ΔH5***GP (*Fluc*-HCV IRES deletion mutant; Fig. 8A) in BHK cells. Again, these di-cistronic RNAs had a non-capped 5’-end (pppA-41 of non-viral nt) with the propensity to form a stable hairpin (see Fig. 3A) to avoid high-level background of *Fluc* expression.

Individual transfection of these di-cistronic transcripts into the BHK cells revealed that upstream expression of *Fluc* was very weak in all cases (Fig. 8B). Expression of *Gluc*, however, was strong under the control of the intergenic ZIKV or DENV 5’-UTRs (Fig. 8C). Compared with the inactive control RNA F***ΔH5***GP the *Gluc/Fluc* ratio indicated an approximately 30-fold higher activity of the F***Z5***G***Z3*** mRNA in the intergenic spacer region (see Fig. S6). Our qRT-PCR results showed that the difference in translational efficiencies is not due to RNA instability of the reporter RNAs isolated from the transfected cells (Fig. S8).

### Silencing of the Xrn-1 gene leads to increased expression of reporter genes with our reporter mRNAs

Throughout the experiments reported here we observed that our mRNAs, if 5’-terminated with pppAN…, were considerably less active in promoting translation of the reporter gene than mRNA terminated with 5’-GpppAN. We thought that this may be due to the function of Xrn-1, a key component in the major 5’-to-3’ mRNA decay in host cytosols (23). We hypothesized that reducing the function of Xrn-1 (“silencing”) may lead to an increase of the signals of our reporter RNAs, including mRNAs 5’ terminated with pppAN.

Silencing of the Xrn-1 gene was achieved by transfection of different amounts of Xrn-1 specific siRNAs (Santa Cruz Biotech, TX), together with an irrelevant control siRNA (IR), into A549 cells (derived from human alveolar basal epithelial cells (24)). Transfection of the cells with siRNAs was accomplished by lipofectamine RNAiMAX (Invitrogen) protocol. Compared to control siRNA (IR)-treated cultures, 10 pmol and 30 pmol Xrn-1-specific siRNA effectively reduced the expression level of Xrn-1 protein at day 1 and day 2 post transfection, respectively. In our experiments with reporter mRNAs D5GD3 and Z5GZ3 we used 30 pmol of Xrn-1 siRNA, which prevented the synthesis of Xrn-1 protein after one day of treatment (Fig. 9A) (see Materials and Methods).

**Fig. 9.**
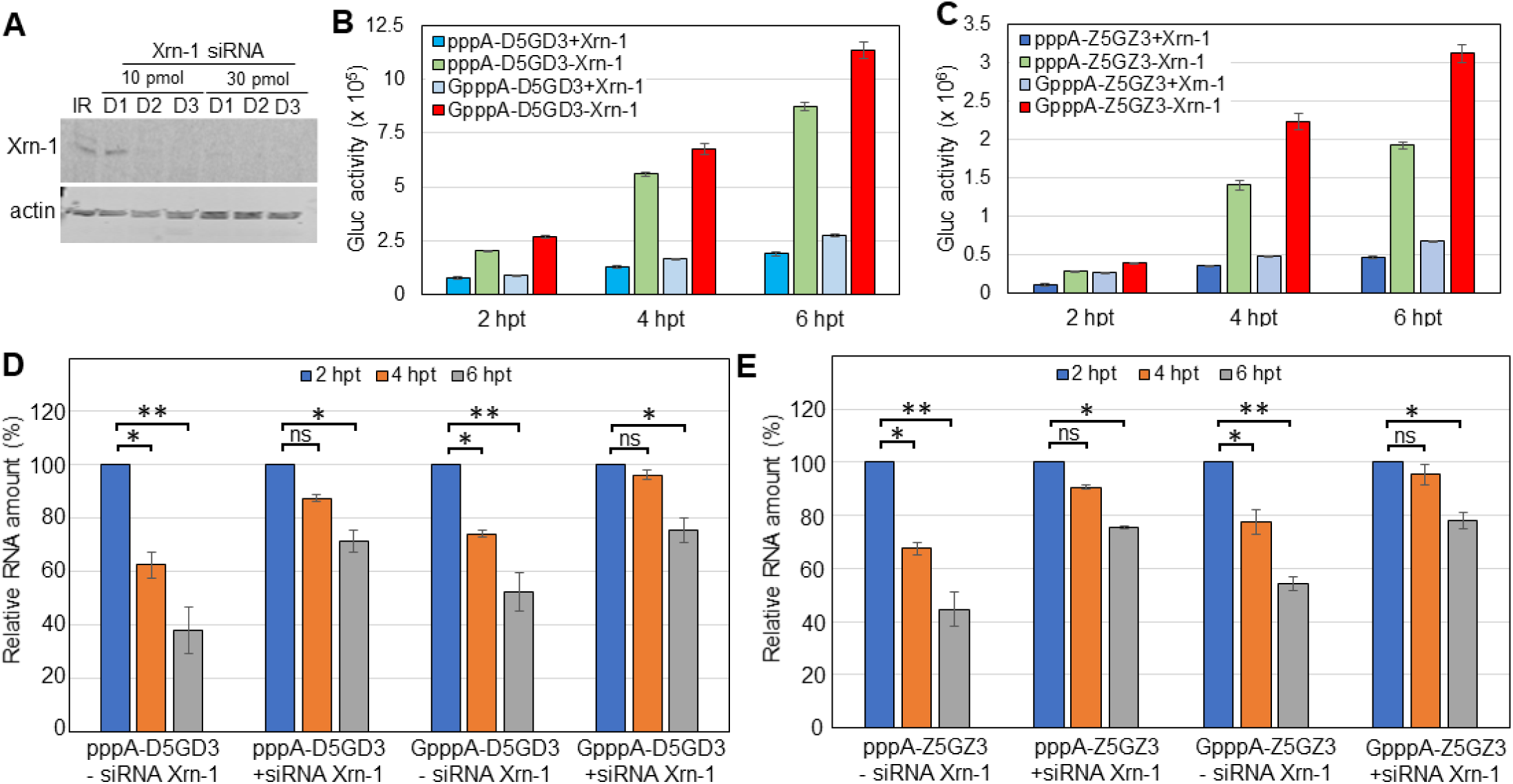
Translation of mono-cistronic reporter RNAs containing the DENV2/ZIKV 5’-UTRs with different 5’-modifications in A549 cells treated with Xrn-1 siRNA. **(A)** Xrn-1-specific siRNA was transfected onto A549 cells with control irrelevant siRNA (IR). Total protein was harvested, and Xrn-1 levels were analyzed by immuno-blotting assay. Actin served as a loading control. **(B, C)** A549 cells treated with (-Xrn-1) and w/o (+Xrn-1) siRNAs were transfected with two types of reporter RNAs, ***D5***G***D3* (B)** and ***Z5***G***Z3* (C)**. *Gluc* activities were measured at different time-points post transfection as indicated. Means of three independent experiments are plotted ± SEM. **(D, E)** RNAs, extracted from ***D5***G***D3* (D)**- and ***Z5***G***Z3* (E)**-transfected A549 cells at different time-points post transfection, were subject to qRT-PCR analysis generating 5’-end fragments. Each reaction product was finally normalized to GAPDH and presented as the fold change relative to input RNAs. The amount of RNAs at 2 hpt was used as time 0 post transfection and artificially set to 1. The means of three independent experiments are plotted ± SEM. A significant difference by two-tailed t-tests compared to control RNA (at 2 hpt) is indicated by * (P<0.05) and ** (P<0.01), respectively. ns, not significant.

All ***D5***G***D3*** and ***Z5***G***Z3*** reporter RNAs, irrespective of their 5’-termini (non-capped pppA- or GpppA-terminated), produced 2-3 fold higher expression levels of *Gluc* in Xrn-1-silenced A549 cells (Figs. 9B and 9C). This result supports our hypothesis that Xrn-1 silencing may have resulted in the protection of mRNAs and led to increased reporter gene signals. Parallel experiments to test mRNA levels by qRT-PCR seemed to support this conclusion as *Gluc* expression co-varied with mRNA stability, and that stability co-varied with the levels of Xrn-1. However, it appeared that GpppA-terminated mRNAs yielded strong *Gluc* signals regardless of the extent of Xrn-1 silencing. We cautiously explain this result with the hypothesis that GpppA-termini may protect against 5’ to 3’ degradation. Clearly, this protection would not be available for RNAs terminated with pppAN-. We note that at the early stage of transfection all mRNAs (pppA- or GpppA-terminated) were relatively stable whereas at wt level of Xrn-1 the relative amount of all reporter mRNAs was reduced very rapidly after 6 hrs post transfection (Figs. 9D and 9E). Among those mRNAs, the non-capped pppA-RNA level reduced the most 6-8 hrs post transfection (Fig. S9). Similarly, the qRT-PCR derived from the non-capped di-cistronic mRNAs F***D5***G***D3*** and F***Z5***G***Z3*** produced almost the same results (data not shown).

## Discussion

The surprising infectivity of purified uncapped RNA transcripts of DENV or ZIKV cDNA clones in mammalian (BHK, Vero) or insect (C6/36) tissue culture cells (Figs. 1 and 6) can be explained by several hypotheses. These include initiation of translation of a small number of intact uncapped viral genomes that allows the formation and amplification of intact replication complexes (25). A second hypothesis draws on reports of cytoplasmic capping of the uncapped RNA molecules post transfection (26). Cytoplasmic capping occurs predominantly on RNA fragments (not necessarily on mRNA fragments) and the selection process for cytoplasmic capping is poorly understood (26). In Fig. 3C we show that the *Fluc* reporter gene still yields very poor translation after incubation of transcripts in BHK cells for as long as 7 hrs, a result which may argue against cytoplasmic capping. However, considering the large number of genome transcripts used in the infection (Fig. S1), a rare and fortuitous progression of uncapped transcripts to repaired, capped genomes cannot be excluded.

An alternative hypothesis as to how the non-capped genomes initiated translation and replication is the function of the 5’-UTR as a minute IRES element. The viral polyprotein, once it has been synthesized under the control of the 5’-UTR, will be proteolytically processed and enzyme functions will emerge that not only catalyze genome synthesis but that will also cap the newly synthesized viral RNAs to produce authentic viral genomes. Intracellular repair of modified 5’ termini of viral transcripts has been observed before (27). For example, poliovirus full-length transcripts with the false 5’ end pppGGUUA had a specific infectivity ∼100 fold lower than that of virion RNA. On transfection of pppGGUUA-terminated transcripts into HeLa cells, newly repaired, authentic virion RNAs (VPg-UUA) emerged in the course of replication.

The experiments with uncapped mono-cistronic mRNAs showed decisively that the DENV 5’-UTR was sufficient to initiate translation of the reporter *Gluc*. Moreover, neither the presence of the DENV 3’-UTR, nor the presence of a polyadenylated 3’ end was essential for, or did interfere with, *Gluc* synthesis. However, the DENV 3’-UTR at the 3’ end of the ***D5***G***D3*** mRNAs stimulated *Gluc* to some extent (Figs. 2B and 2C), a result for which we have no explanation.

All mono-cistronic mRNAs carrying the DENV or the ZIKV 5’-UTRs terminated with an uncapped 5’ pppAN- were weak in inducing *Gluc* translation when compared with 5’ GpppA-mono-cistronic mRNAs. We now have evidence that this deficiency is related to RNA stability. In BHK cells the mRNAs are possibly degraded 5’→3’ by the Xrn-1 as shown by silencing the Xrn-1 gene (Figs. 9D and 9E). Interestingly, mRNAs 5’-terminated with uncapped GpppAN are considerably more resistant to degradation than mRNAs with pppAN-termini regardless of the degree of Xrn-1 silencing (5’-terminal blocking of 5’→3’ exonucleolytic degradation?). We are not sure at present whether other factors play a role in the weak induction of pppAN-terminated mRNAs.

The standard test to detect IRES activity in an RNA sequence is the di-cistronic reporter assay (1, 2). Accordingly, we cloned the DENV or ZIKV 5’-UTRs into the intergenic region of di-cistronic mRNAs (*Fluc*-x-*Gluc*) followed by transfection into the mammalian BHK cells. We note that the 5’-terminal sequence preceding *Fluc* ORF in our di-cistronic mRNAs was an uncapped sequence capable of forming a stable RNA stem-loop (Fig. 3A). Thus, it is unlikely that this 5’ terminus supports false initiation of translation and high *Fluc* background signals. Based on the observed strong signals of the downstream *Gluc* reporter we conclude that the short 5’-terminal UTRs of DENV or ZIKV are RNAs with IRES competence for internal ribosomal entry. Similar results were obtained with other di-cistronic mRNAs that carried either the IRESs of mammalian viruses (HCV or EMCV; ∼350 to ∼450 nt each), or the small intragenic IRES of CrPV (189 nt) (3) in the intergenic regions. We believe that our results with di-cistronic mRNA that carry the IRESs of HCV, EMCV, or CrPV in the intergenic region, serve as important controls.

Repeating these experiments in C6/36 cells, however, yielded a surprise: neither the DENV nor the ZIKV 5’-UTR, when placed into the intergenic region of di-cistronic mRNAs, expressed IRES function (activation of *Gluc* synthesis) in insect cells. Similarly, when placed into the intergenic region of di-cistronic mRNA the IRESs from other mammalian viruses (HCV and EMCV) were also inactive in C6/36 cells (Fig. 5). On the other hand, a strong *Gluc* signal was obtained when the small intergenic IRES of CrPV (189 nt) was placed between *Fluc* and *Gluc* (Fig. 5C, F***Cr***GP RNA). This experiment supports the validity of the di-cistronic mRNA experiment.

The IRESs of HCV, a *Hepacivirus* of *Flaviviridae*, and of EMCV, a cardiovirus, are also inactive in insect cells (Fig. 5) as had been shown by others (21, 22). The reason for the lack of activity of the flavivirus 5’-UTRs in di-cistronic mRNA in mosquito cells could point to a lack of specific cellular translation factors, or of ITAFs, or RNA misfolding in insect cells. However, since both DENV and ZIKV are arboviruses one would have expected an IRES-like function of their 5’-UTRs in insect cells in both mono- and in di-cistronic mRNAs.

An unorthodox interpretation of these unexpected results is that the IRES competence of DENV or ZIKV 5’-UTRs is a remnant from ancestor viruses of *Flaviviridae* that now belong to the genera *Hepacivirus, Pestivirus*, or *Pegivirus*, e.g. viruses that may never have replicated in arthropods. This could explain the inactivity of the HCV IRES and of the DENV and ZIKV 5’-UTRs when placed into a di-cistronic cassette and tested in insect cells.

IRES elements are amongst the most mysterious RNA structures in eukaryotic molecular biology. As the name suggests (28), an IRES allows initiation of translation of a eukaryotic mRNA independent of the nature of the 5’ terminus of the mRNA. Although discovered 30 years ago, the origin of IRES elements during evolution is unknown. Inexplicably, IRESs are defined by function rather than by size or structure (29). Since single stranded RNAs can fold innumerable ways to innumerable structures one can envision that one or more of these structures can lead to complexes between the ribosomal subunits, canonical translation factors and, possibly, cellular proteins (ITAFs). In a large screen, many of those complexes may lead to translation initiation without clear biological relevance. Remarkably, Wellensiek et al, (30) searching for cap-independent translation enhancing elements, found >12,000 of such entities in the human genome. Most of these elements still await further characterization (30). These studies have been extended recently by Weingarten-Gabbay et al., (31) who reported the existence and function of thousands of human and viral sequences, expressing specific signatures of cap-independent translation activity. We stress that the function of an IRES is expressed most efficiently in cells (or their extracts), which are related to the natural habitat of the virus to which the IRES belongs and, thus, are likely to provide proteins required for the virus-specific IRES function. For example, translating poliovirus genomes in rabbit reticulocyte lysates revealed an IRES in the C-terminal part of its polyprotein, which turned out to be an artifact (32). Adding protein factors from HeLa cells to the translation extract (33), or translating the poliovirus genome in HeLa extracts altogether (25), completely eliminated this artifact. The cellular proteins necessary for the IRES functions of the DENV and ZIKV 5’-UTRs are not yet known.

Regardless of whether IRESs are large (∼450 nt; poliovirus, encephalomyocarditis virus) or small (e.g. 189 nt; intergenic region IRES in dicistroviruses) they are RNA segments that require the cooperation of cellular protein factors for function. This is in contrast to prokaryotic systems. For example, ORFs in multi-cistronic mRNAs of *E. coli* are preceded by the *Shine-Dalgarno* sequence (S/D sequence (5’**-*AGGAGG*-**3’) (34, 35) that alone functions as an internal ribosomal entry site by binding to a complementary sequence at the 3’ end of the ribosomal 16S rRNA (5’**-**GATCA***CCUCCU***UA**-**3’). The discovery of eukaryotic IRESs prompted also a search for RNA sequences in the IRES that could function like S/D sequences. Early work concentrated on oligopyrimidine segments in picornavirus IRESs that, however, were found to serve in specific combinations with other segments of the IRES as “ribosomal landing pads” (36, 37). An exceptional case is a 9 nt-long sequence (5’-***CCGGCGGGU***-3’) found in an IRES of the mRNA encoding the *Gtx* homeodomain protein, which activates translation by base pairing to 18S ribosomal RNA (38, 39). We have searched in vain in DENV and ZIKV 5’UTRs for sequences complimentary to 18S rRNAs.

Currently, we cannot answer the question why the 5’ UTRs of DENV and ZIKV, presumably of related flaviviruses [West Nile virus (WNV), yellow fever virus (YFV), Japanese encephalitis virus (JEV), etc.], harbor an IRES function. We speculate that a role of the minute flavivirus IRES could emerge during virus infection when cellular mechanisms scale down cap-dependent translation to reduce proliferation of the invading pathogen (40). Moreover, it is possible that flaviviruses continue their translation through the metaphase of cells, when cap-dependent translation is reduced (41).

The work described here raises many questions about flavivirus genetics and control of proliferation. As we have implied above, another intriguing puzzle to be solved is at what stage in evolution one group of the *Flaviviridae* decided to restrict regulation of translation to a specific mechanism at the expense of another mechanism, and why?

## Materials and Methods

### Cell cultures

BHK (Baby Hamster Kidney) cells, A549 cells (adenocarcinomic human alveolar basal epithelial cells), and Vero cells were maintained in Dulbecco’s modified eagle medium (DMEM) supplemented with 10% Fetal bovine serum (FBS) (HyClone Laboratories, Logan, UT), 100 U/ml of penicillin, and 100 µg/ml of streptomycin (Invitrogen). All cells were grown at 37°C in a 5% CO_2_ incubator. Mosquito C6/36 (ATCC, Bethesda, MD), an *Aedes albopictus* cell line, was cultured in minimal essential medium (MEM, Gibco BRL) supplemented with 1/100 non-essential amino acids (NEAA, Gibco BRL), 10% FBS, penicillin G (100 U/ml) and streptomycin (100 mg/ml) at 28°C in a 5% CO_2_ incubator.

### Plasmids

(i) p***D5***G***D3***, p**D5**GP, pGP, p***H5***G***H3*** mono-cistronic constructs. p***D5***G***D3*** or p***D5***GP, containing the entire DENV type 2 5’-UTR sequences (96 nts), *Gluc* encoding sequence (New England Biolabs, NEB) followed by the DENV2 3’-UTR or unrelated poliovirus 3’-UTR and a poly(A) tail, was constructed by standard cloning strategy. In order to produce robust run-off in vitro transcripts, a modified T7 promoter sequence was place immediately upstream of the authentic DENV2 5’-UTR starting with an Adenosine (10). pGP was modified from a commercially available plasmid pCMV-Gluc (NEB), in which contains a set of cellular 5’-noncoding sequences (41 nts) and Poliovirus 3’-UTR followed by a poly(A) tail in replacement of its original 3’-UTR sequences (Fig. 2A). p***H5***G***H3*** is a mono-cistronic construct, in which the *Gluc* ORF is flanked by an HCV IRES at the 5’ end and the HCV 3’-UTR at the 3’ end, respectively (Fig. 4).

(ii) Di-cistronic reporter constructs. All di-cistronic reporter constructs: pF***D53***GP, pF***D35***GP, pF***E***GP, pF***D3***GP, pF***D5***G***D3***, pF***Z5***G***Z3***, pF***D5***GP, pF***Cr***GP, pF***H5***GP, and pF***ΔH5***GP plasmids contain the T7 RNA polymerase promoter linked to a cellular 5’-noncoding sequences including a stable stem-loop (41 nucleotides, nts, Fig. 3A) fused to the *Fluc* encoding sequences followed by two termination codons. For pF***H5***GP/pF***ΔH5***GP di-cistronic constructs, the second cistron contains an active HCV IRES element, or an inactivated HCV IRES element (in which the stem-loop III, from nucleotide 133 to 290, within the 5’-UTR is entirely deleted), including the 5’ beginning part of core protein encoding sequences followed by a cellular ubiquitin sequences, then fused with *Gluc* coding sequences ending with a stop codon. The presence of a cellular ubiquitin sequences is to produce 5’-authentic end of downstream *Gluc* coding sequences. For pF***D53***GP and pF***D35***GP di-cistronic constructs, a cap-dependent cellular 5’-UTR fused to the *Fluc* gene sequence as a first cistron, followed by two stop codons to prevent read-through. The second cistron consists of sequences of the DENV 5’- and 3’-UTR in either 5’-3’ (a Kozak context was inserted between the 3’-UTR and *Gluc* encoding sequence), or 3’-5’ orientation, then directly fused to the *Gluc* reporter gene-encoding sequences and finally an unrelated 3’ non-coding sequence followed by a poly(A) tail. For pF***D5***G***D3*** and pF***D5***GP di-cistronic constructs, translation of the *Gluc* reporter gene is initiated by a DENV 5’-UTR sequence (96 nts) in the presence of DENV 3’-UTR sequence (451 nts) only (F***D5***G***D3***) and an unrelated 3’-noncoding sequence (derived from poliovirus genome sequence) followed by a poly (A) tail (F***D5***GP), respectively. For di-cistronic constructs pF***E***GP, pF***D3***GP, and F***Cr***GP, the general structure is the same as pF***D5***GP except for the internal initiation of translation was directed by an EMCV IRES (F***E***GP), a DENV 3’-UTR sequence (F***D3***GP), and CrPV IGR IRES (189 nts, F***Cr***GP), respectively. For pF***Z5***G***Z3*** construct, translation of the *Gluc* reporter gene is initiated by a ZIKV 5’-UTR sequence (107 nts) in the presence of ZIKV 3’-UTR sequence (428 nts) at the very 3’-end.

### Flaviviral infectious clones

The infectious dengue virus serotype 2 clone DENV2^syn^ was synthesized and assembled in our lab (42). An infectious cDNA ZIKV clone (strain FSS13025, GenBank number KU955593.1) developed by Dr. Pei-Yong Shi’s laboratory at University of Texas Medical Branch (UTMB), Galveston, TX was kindly gifted to us (43).

### *In vitro* transcription and RNA transfection in both mammalian and insect cells

Plasmids were purified by standard Qiagen procedure. The templates for *in vitro*-transcription of the reporter RNAs were generated by either PCR (for RNA ending with DENV 3’-UTR) or cleavage linearization (for RNA ending with regular poly-A tail) to obtain run-off transcripts with the precise 3’-ends, followed by phenol/chloroform extraction and ethanol precipitation. About 1 µg of purified RNA template was transcribed by T7 RNA polymerase (NEB) and the integrity of the RNAs was examined by agarose gel electrophoresis. RNAs were purified by Qiagen RNeasy kit and re-dissolved in double distilled H_2_O. The concentration of RNAs were determined by a spectrophotometer Nanodrop-1000 (Thermo Fisher Scientific). Between 0.4 and 1 µg of transcript RNA was used to transfect BHK or C6/36 cells on a 12-plate by Lipofectamine^®^2000 protocol as described previously (44).

Transfected BHK and C6/36 cells were incubated in 5% FBS-DMEM at 37°C (for BHK or Vero) and 5% FBS-MEM at 28°C (for C6/36), respectively. BHK or Vero cells were checked daily for cytopathic effect (CPE). When CPE was detected, infection medium was harvested and fresh 5% FBS-DMEM was added for further incubation till day 9 post transfection. For transfected C6/36 cells, the medium was harvested at day 5 post transfection and fresh 5% FBS-MEM was added for further incubation. The supernatant containing the viral particles was aliquoted and stored at −80°C for further passaging or infection. Viral titers were determined by a modified plaque assay on BHK or Vero cell monolayers using agarose overlay as previously described (45).

### siRNAs and siRNA Assay

Xrn-1 siRNA was purchased form Santa Cruz Biotechnology, Inc., Dallas, TX (Cat #: sc-61811). Another irregular siRNA (si IR) was designed as a duplex with UU 3’ overhangs with the following sequence: 5’-AAGGACUUCCAGAAGAACAUC-3’ taken from (46), and was synthesized by Eurofins Genomics LLC (Louisville, KY). Different amounts (10-30 pmol) of RNA duplexes were transfected with Lipofectamine RNAiMAX reagent (Thermo Fisher Scientific) onto A549 cells at the confluence of 60%-70%. Cells were maintained in standard conditions. At different time-points (up to 3 days) post transfection, cell lysates were prepared and eventually subjected to SDS-PAGE. The silencing effect was examined by western blot with specific anti-Xrn-1 antibody (Boster Biological Technology, Pleasanton, CA).

### RNA isolation from transfected/infected cells and quantitative RT-PCR

70-80% confluent BHK or C6/36 cells, seeded on 12-well plate or 35-mm-diameter plate, were transfected with reporter RNAs or infected with virus supernatants at ∼1 PFU per cell. At different time-points post transfection (or when CPE was observed during infection of BHK cells), total RNA was extracted from either transfected cells or 200 µl of viral supernatant with 800 µl *Trizol*^®^ reagent (Invitrogen), according to the manufacturer’s instruction. Quantitative RT-PCR was performed and analyzed according to the protocols of StepOnePlus™ Real-Time PCR System (Thermo Fisher Scientific) with 2× SYBR green master mix (Quanta Biosciences). PCR was performed with primers located on 5’- and 3’-end of ORF of individual reporter gene, respectively. Detailed information of oligos was described in Supplementary Materials. Values were normalized to that of GAPDH, which was amplified with primer pairs of 5’-ATGGCCCCTCCGGGAAACTG and 5’-ACGGAAGGCCATGCCAGTG.

### Assay of luciferase activities

At the indicated time-points, 10-20 µl of the culture medium was taken for the measurement of *Gluc* activity. To assay cell lysates for *Fluc* expression, transfected cells growing in 12-well plates were carefully washed twice with warm phosphate-buffered saline (PBS) followed by treatment with 120 µl 1× Passive Lysis Buffer (PLB, Promega) by shaking for 10-15 min at room temperature. The supernatant was collected and cleared by centrifugation at 4°C. Luciferase activity was determined in a luminometer (Optocomp I) by the addition of ∼18 µl of sample to 6-7 µl diluted coelenterazine (for *Gluc*, NEB), or ∼20 µl of sample to 15-18 µl *Fluc* substrate (Promega).

### Statistical analysis

Calculation of the mean and standard deviation (SD) were performed by Microsoft Office 365 (Microsoft Corporation, WA). A statistically significant differences were compared by a two-tailed, unpaired, and unequal variant Student’s t-test.

## Acknowledgments

We are indebted for suggestions from and discussions of this work with A. Paul, J. Cello, and O. Gorbatsevych, and we thank Dr. Pei-Yong Shi (Galveston, TX) for a gift of the ZIKV infectious cDNA clone, and Dr. Patrick Hearing and Dr. David Thanassi for helpful discussion about statistical analysis. We gratefully acknowledge A. Wimmer for editing our manuscript. This work was supported in parts by grants from the National Institute of Health RO1AI07521901 (E.W.) and 1R21AI126048-01 (Y.S.).

## Supplemental Materials

### Primers for qRT-PCR

PCR fragments were amplified in qRT-PCR with the following primers: for 5’-end fragments (129 bps) from mono-cistronic RNAs: 5’-AGTTCTGTTTGCCCTGATCTGC and 5’-AACTTCCCGCGGTCAGCATC; for 5’-end fragments (129 bps) from di-cistronic RNAs: 5’-AAGGCCCGGCGCCATTCTATC and 5’-ATGTTCACCTCGATATGTGCTC; for 3’-end fragments (136 bps) from both mono- and di-cistronic RNAs: 5’-ATCTGTGTGTGGACTGCACAAC and 5’-TTGATCTTGTCCACCTGGCC; for GAPDH fragment (137 bps): 5’-ATGGCCCCTCCGGGAAACTG and 5’-ACGGAAGGCCATGCCAGTG.

**Fig. S1.**
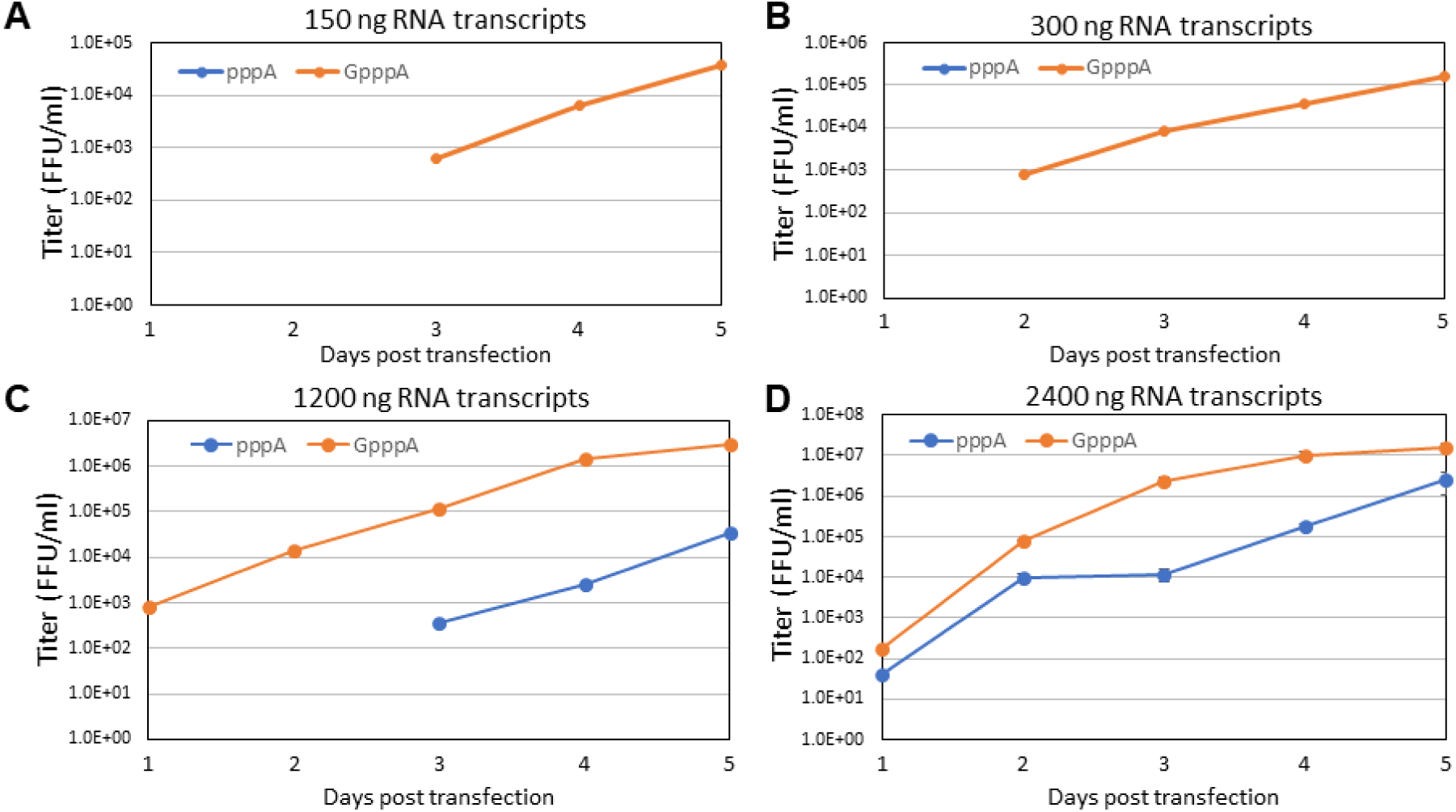
Infectious assays of different amounts of non-capped pppA- and GpppA-terminated DENV transcript RNAs in mammalian cells. Equal amount of two types of DENV2 RNA transcripts were transfected onto BHK monolayers. Culture media were collected daily (up to day 5) and viral titers were measured by focus forming assay in Vero cells.

**Fig. S2.**
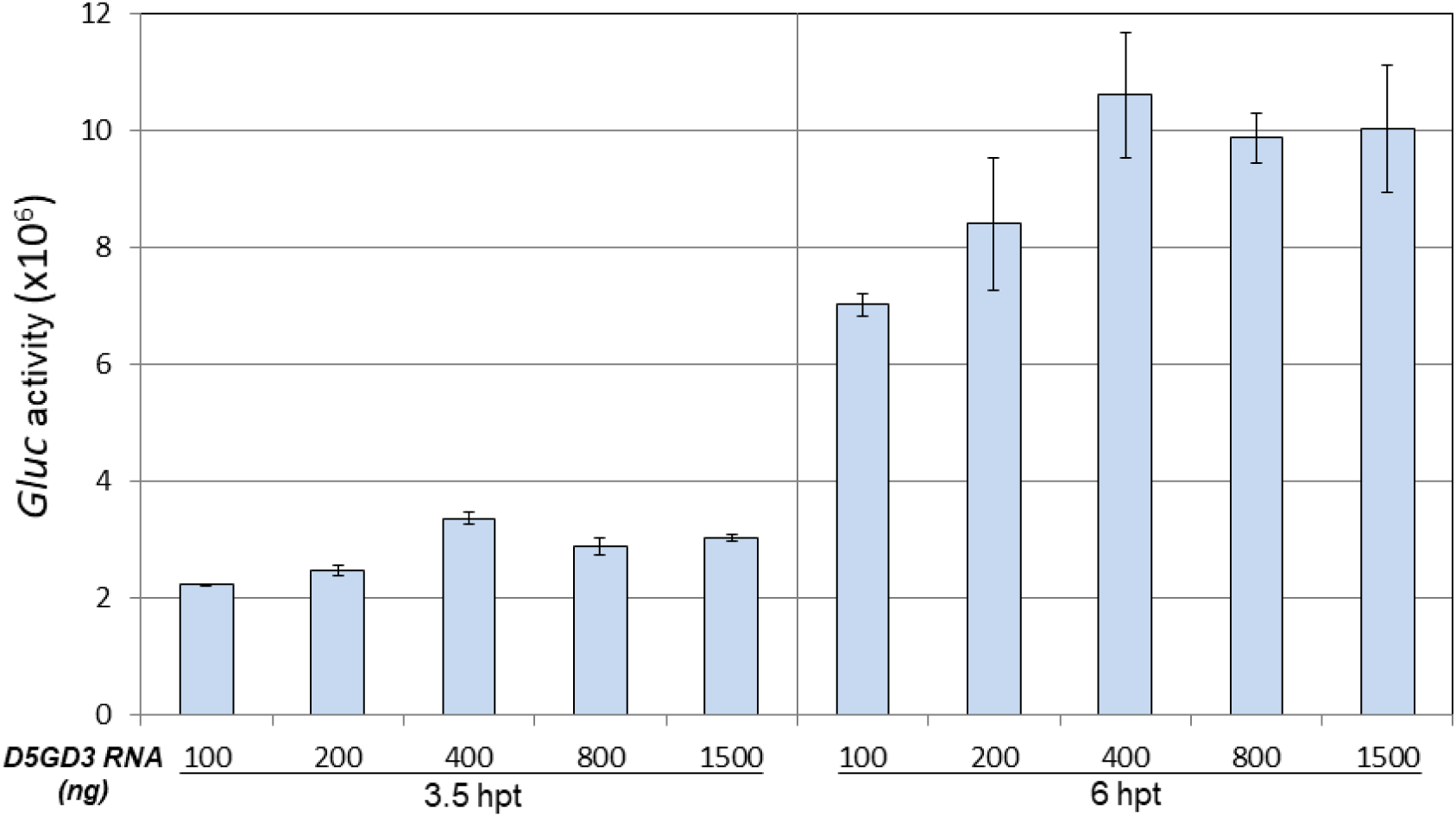
Translational efficiency directed by different amounts of *D5*G*D3* NAs in BHK cells. Different amounts of RNA were transfected into BHK monolayer cells seeded on a 12-well plate. The *Gluc* activity was measured at different time-points post transfection as indicated. The means of three independent experiments are plotted ± SEM.

**Fig. S3.**
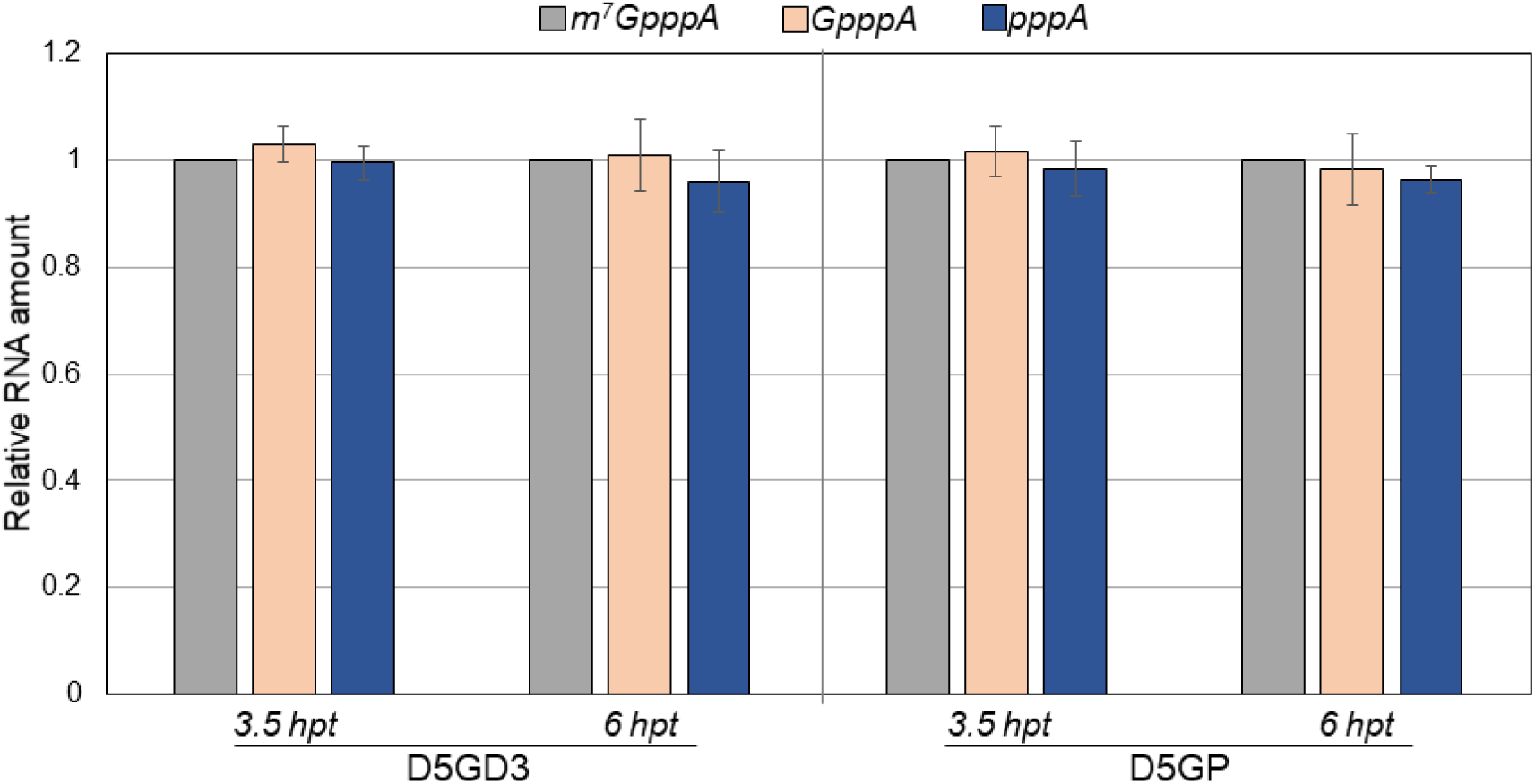
Relative amount of mono-cistronic RNA transcripts in transfected cells analyzed by qRT-PCR. Total RNA was extracted and isolated by *Trizol*^®^ reagent from cells transfected with RNA transcripts at different time-points post transfection. RNA levels were measured by qRT-PCR and normalized by GAPDH. The relative amount of m^7^GpppA RNA was artificially set to 1. The means of three independent experiments are plotted ± SEM. P values showed no significant difference between samples at indicated time points.

**Fig. S4.**
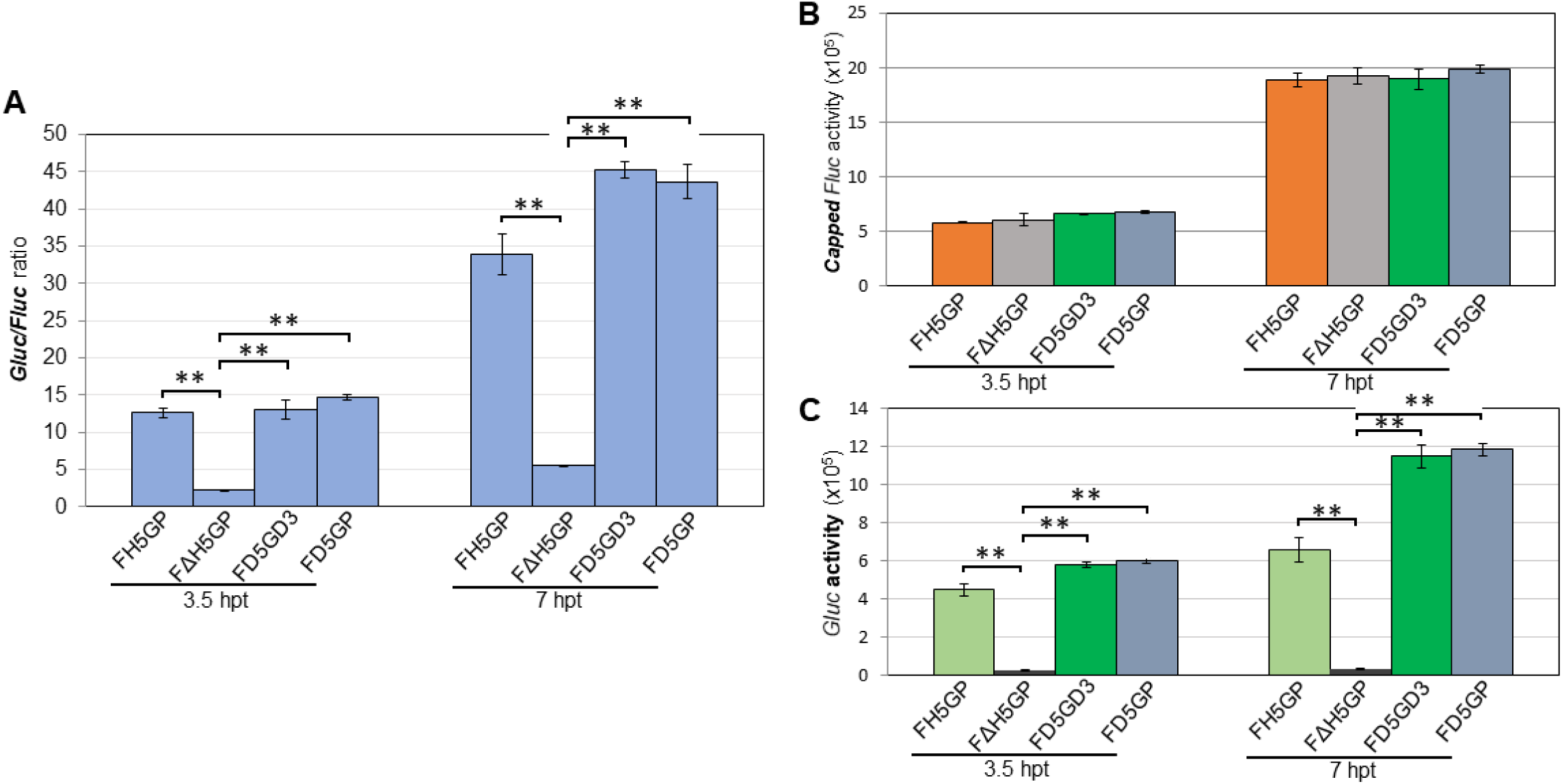
The DENV2 5’-UTR confers IRES activity in di-cistronic mRNAs. (A) Relative *Gluc* expression level normalized by *Fluc* activity, associated with Fig. 3. (B) Translation efficiency of 5’ capped *Fluc* reporter. F***H5***GP and F***ΔH5***GP reporter RNAs were included as positive and negative control, respectively. **(C) *Gluc* expression directed by internal initiation of either the HCV IRES or the DENV2 5’-UTR.** *Fluc* and *Gluc* activities were measured at different time-points post transfection as indicated. The means of four independent experiments are plotted ± SEM. The P values were determined by comparing to the negative control (F***ΔH5***GP) using two-tailed t-tests at the indicated time point. ** indicates P<0.01.

**Fig. S5.**
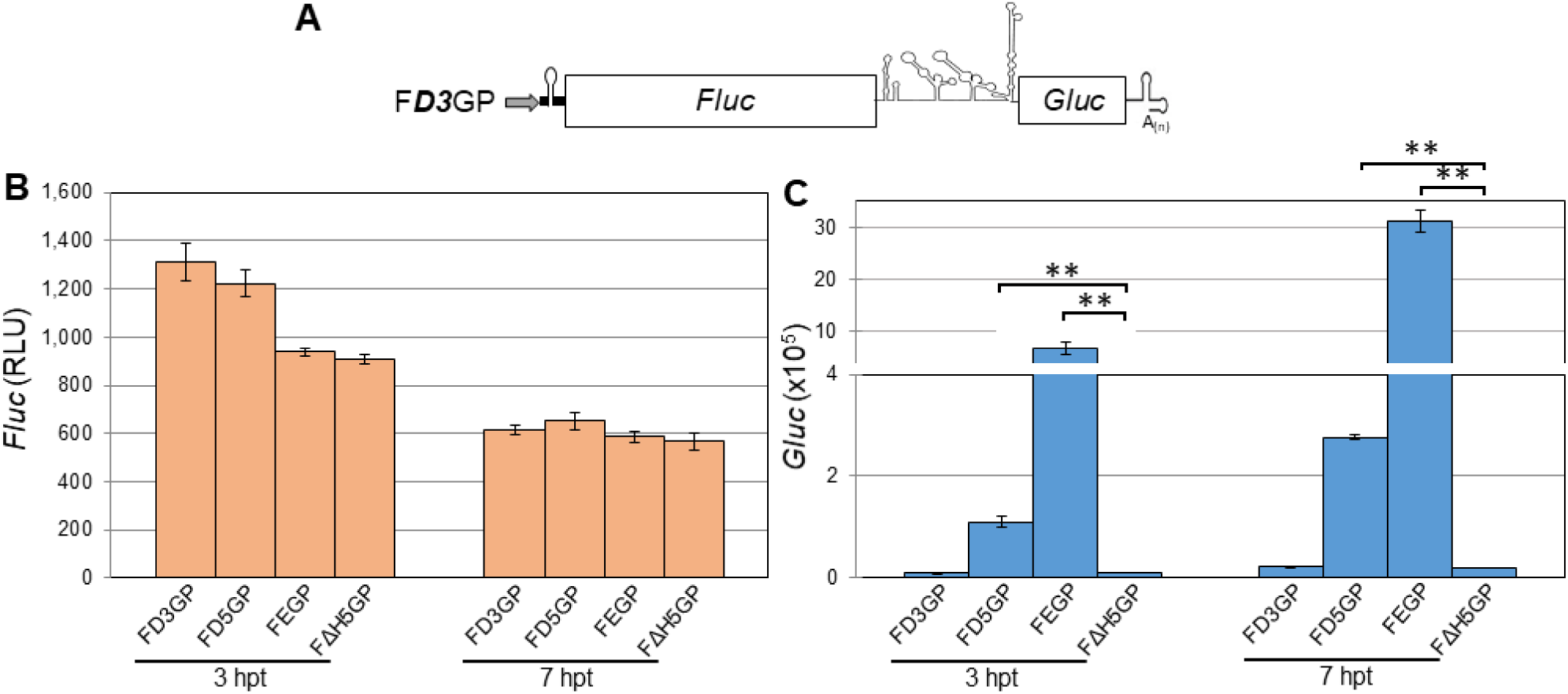
The role of the DENV2 3’-UTR in translation initiation. (A) Schematic diagram of di-cistronic reporter constructs F***D3***GP and F***E***GP. (B) Translation efficiency of 5’ non-capped reporter RNAs. Di-cistronic F***E***GP and F***ΔH5***GP reporter RNAs (shown in Fig. 3B) were also included as positive and negative control, respectively. (C) *Gluc* expression directed by internal initiation of either the HCV IRES or the DENV2 3’-UTR. *Fluc* and *Gluc* activities were measured at different time-points post transfection as indicated. The means of three independent experiments are plotted ± SEM. The P values were determined by comparing to the negative control (F***ΔH5***GP) using two-tailed t-tests at the indicated time point. ** indicates P<0.01.

**Fig. S6.**
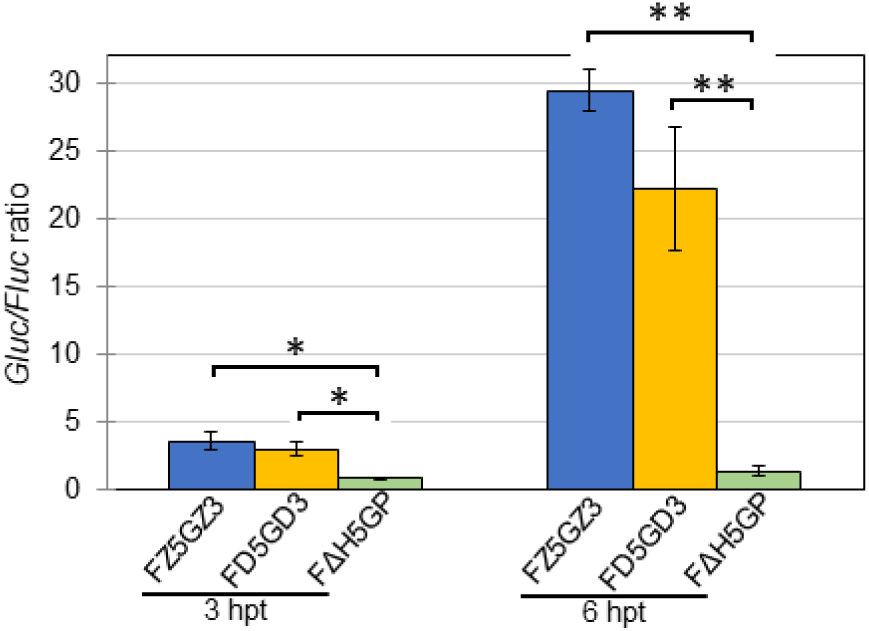
Relative *Gluc* expression level normalized by *Fluc* activity (*Gluc/Fluc*), associated with Fig. 8. The means of four independent experiments are plotted ± SEM. The P values were determined by comparing to the negative control (F***ΔH5***GP) using two-tailed t-tests at the indicated time point. * and ** indicates P<0.05 and P<0.01, respectively.

**Fig. S7.**
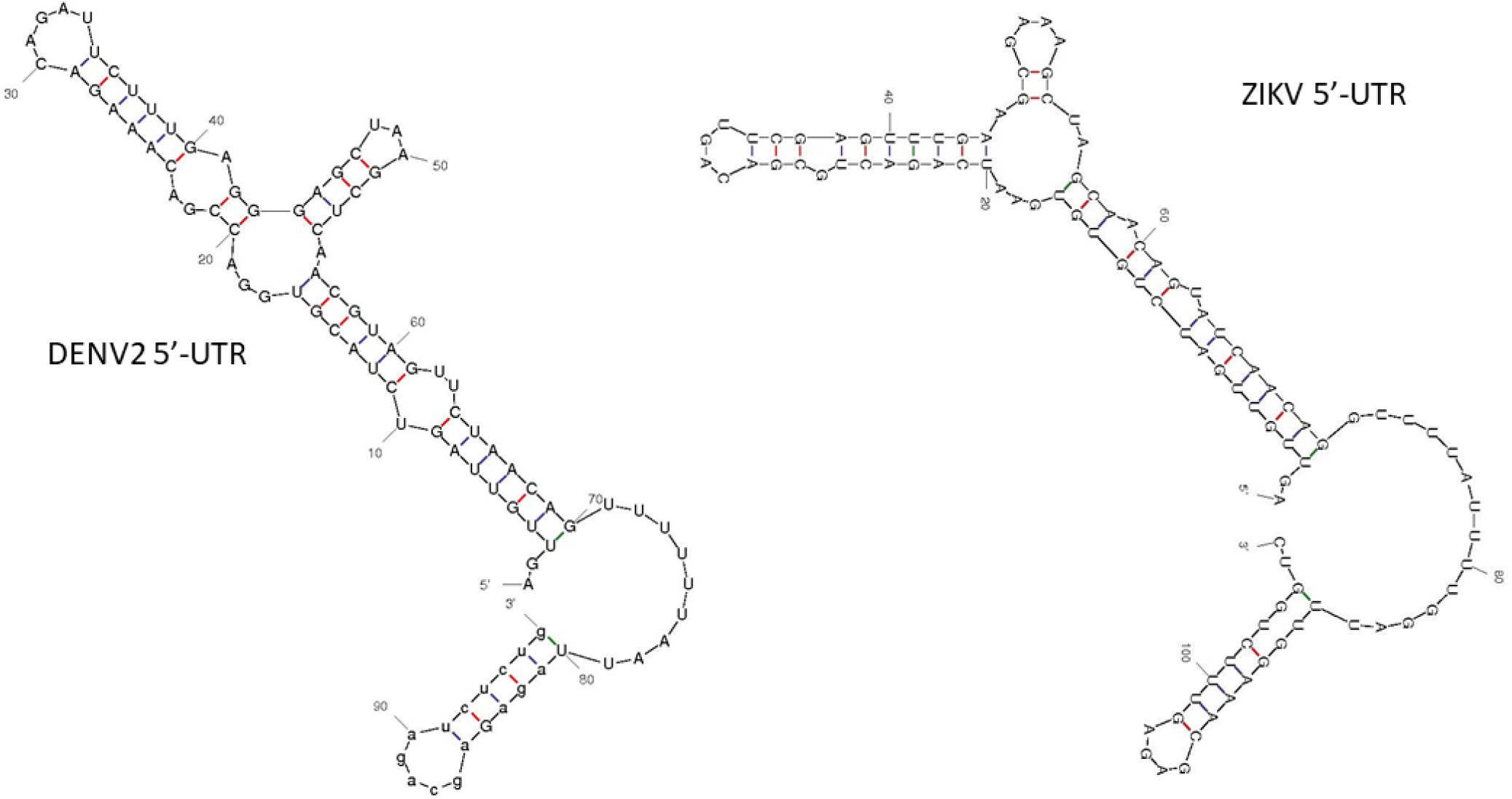
Predicted secondary structure of 5’-UTRs of DENV2 and ZIKV generated by m-fold. DENV2 5’-UTR, initial ΔG = −28.8 kcal/mol; ZIKV 5’-UTR, initial ΔG = −35.90 kcal/mol.

**Fig. S8.**
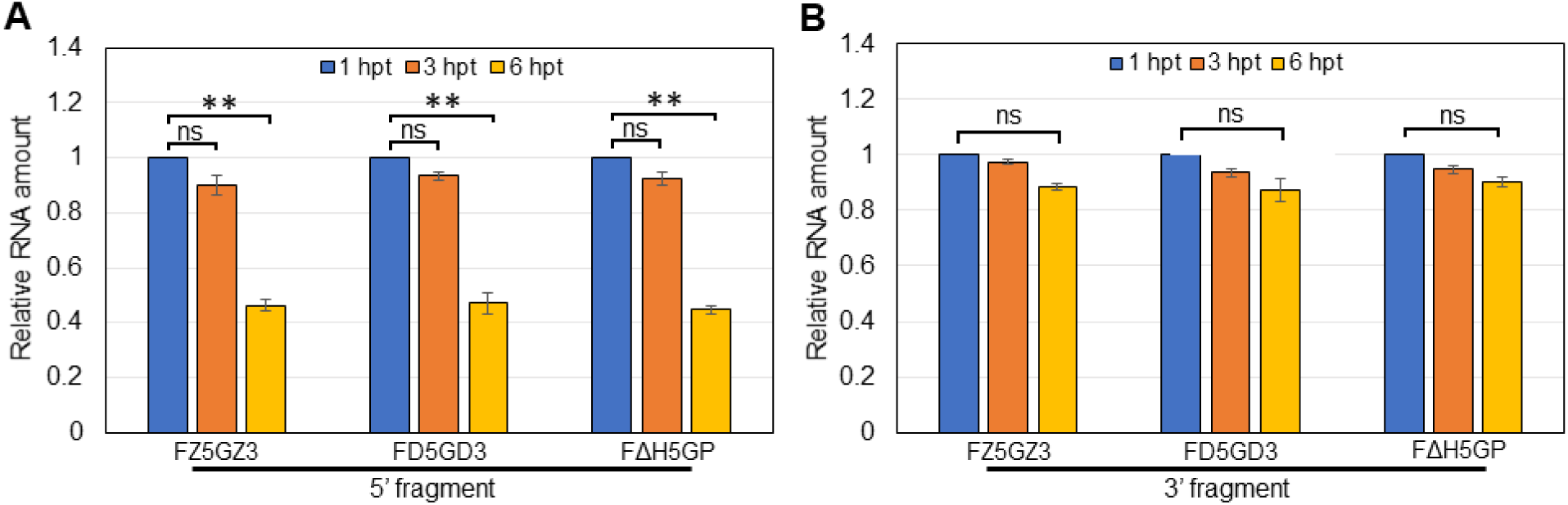
Relative amount of three di-cistronic RNA transcripts in transfected cells analyzed by qRT-PCR. Total RNA was extracted and isolated by *Trizol*^®^ reagent from cells transfected with RNA transcripts at different time-points post transfection. Relative RNA levels were measured by qRT-PCR with either 5’-end oligo primers **(A)** or 3’-end oligo primers **(B)** and calibrated by the RNA amount at 1 hpt (used as time 0 post transfection), respectively. Final percentage of RNAs were normalized by GAPDH. The means of three independent experiments are plotted ± SEM. A significant difference by two-tailed t-tests compared to control RNA (at 1 hpt) is indicated by ** (P<0.01). ns, not significant.

**Fig. S9.**
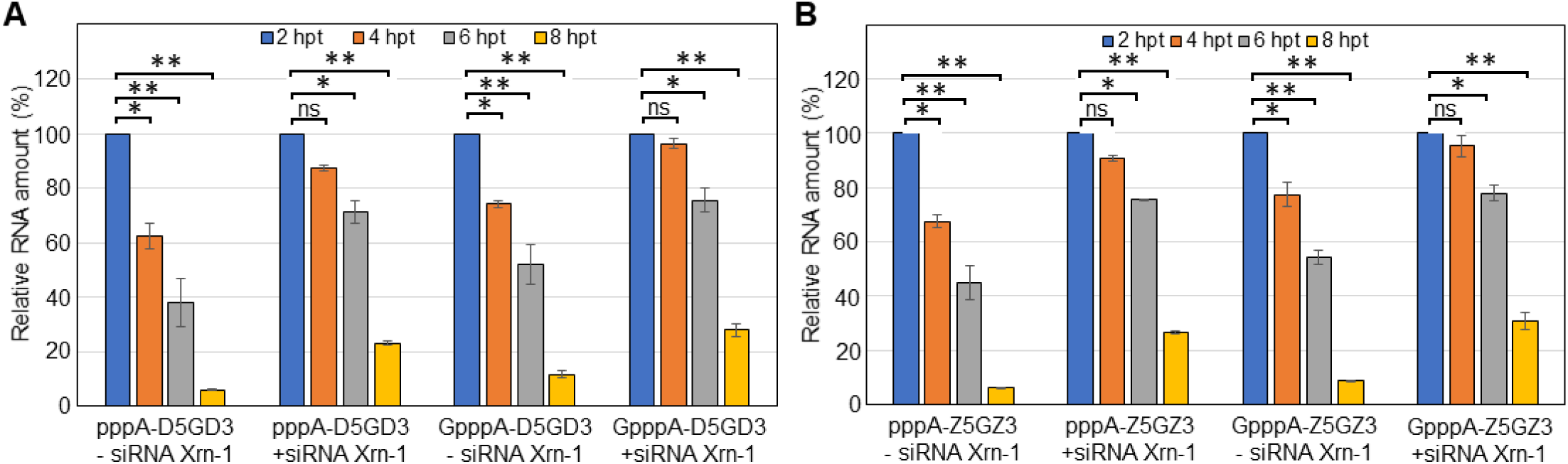
*D5*G*D3*. **(A)** and ***Z5***G***Z3* (B)** RNAs, extracted from transfected A549 cells at different time-points post transfection, were subject to qRT-PCR analysis (5’ end target). Each reaction product was finally normalized to GAPDH and presented as the fold change relative to input RNAs. The amount of RNAs at 2 hpt was used as time 0 post transfection and artificially set to 1. The means of three independent experiments are plotted ± SEM. A significant difference by two-tailed t-tests compared to control RNA (at 2 hpt) is indicated by * (P<0.05) and ** (P<0.01), respectively. ns, not significant.

